# A Sea Change in Microbial Enzymes: Heterogeneous latitudinal and depth-related gradients in bulk water and particle-associated enzymatic activities from 30°S to 59°N in the Pacific Ocean

**DOI:** 10.1101/2020.10.20.346601

**Authors:** John Paul Balmonte, Meinhard Simon, Helge-Ansgar Giebel, Carol Arnosti

**Author notes:** Corresponding author: John Paul Balmonte.

## Abstract

Heterotrophic microbes initiate organic matter degradation using extracellular enzymes. Our understanding of differences in microbial enzymatic capabilities, especially among particle-associated taxa and in the deep ocean, is limited by a paucity of hydrolytic enzyme activity measurements. Here, we measured the activities of a broad range of hydrolytic enzymes (glucosidases, peptidases, polysaccharide hydrolases) in epipelagic to bathypelagic bulk water (non-size fractionated), and on particles (≥ 3 μm) along a 9,800 km latitudinal transect from 30°S in the South Pacific to 59°N in the Bering Sea. Individual enzyme activities showed heterogeneous latitudinal and depth-related patterns, with varying biotic and abiotic correlates. With increasing latitude and decreasing temperature, lower laminarinase activities sharply contrasted with higher leucine aminopeptidase (leu-AMP) and chondroitin sulfate hydrolase activities in bulk water. Endopeptidases (chymotrypsins, trypsins) exhibited patchy spatial patterns, and their activities can exceed rates of the widely-measured exopeptidase, leu-AMP. Compared to bulk water, particle-associated enzymatic profiles featured a greater relative importance of endopeptidases, a broader spectrum of polysaccharide hydrolases, and latitudinal and depth-related trends that paralleled variations in particle fluxes. As water depth increased, enzymatic spectra on particles and in bulk water became narrower, and diverged more from one another. These distinct latitudinal and depth-related gradients of enzymatic activities underscore the biogeochemical consequences of emerging global patterns of microbial community structure and function, from surface to deep waters, and among particle-associated taxa.

## Introduction

Heterotrophic microbial communities play a critical role in the global carbon cycle by transforming and remineralizing up to 50% of marine primary production (Azam and Malfatti, 2007). To initiate these processes, marine heterotrophic microbes secrete hydrolytic enzymes that cleave high molecular weight (HMW) compounds to sizes sufficiently small for bacterial uptake (Weiss et al. 1991). Microbial enzyme activities therefore determine the nature and quantity of compounds available to heterotrophic microbes for biomass incorporation or respiration. Particulate or surface-adsorbed HMW substrates that remain unused in the water column may eventually fuel benthic heterotrophic organisms, or be sequestered in sediments over longer timescales (Arnosti, 2011). However, the specific substrates potentially accessible to microbial communities across vast expanses in the oceans are not well characterized, due in part to sparse enzyme activity measurements, particularly in deep waters and on particles.

Variations in hydrolytic enzyme activities in surface waters indicate that there are spatially distinct microbial capabilities to initiate organic matter remineralization. Prominent differences in enzyme activities emerge especially along latitudinal gradients (Arnosti et al. 2011; Christian and Karl, 1995). Leucine aminopeptidase (leu-AMP) and β-glucosidase exhibit contrasting trends with temperature and latitude in disparate locations (Christian and Karl, 1995). The rates and range of specific polysaccharide hydrolase activities peak in warm temperate waters and decrease towards polar regions (Arnosti et al. 2011). With few exceptions (Ladau et al. 2013), latitudinal trends for enzyme activities mimic those observed for microbial community composition, diversity, and metabolic potentials (Fuhrman et al. 2008; Hewson et al. 2009; Pommier et al. 2007; Wietz et al. 2010; Ibarbalz et al. 2019). These findings suggest the importance of variations in microbial community structure in shaping differences in microbial enzymatic capabilities (Arnosti et al. 2011, Balmonte et al. 2019), and overall metabolic potentials (Raes et al. 2011; Sunagawa et al. 2015; Salazar et al. 2019). However, environmental conditions can be more proximate drivers of microbial function (Louca et al. 2016; Raes et al. 2011; Raes et al. 2018; Sunagawa et al. 2015). The extent to which these parameters correlate with enzyme activities across large spatial scales remains to be tested.

Differences in microbial community composition and environmental conditions may lead to changes in enzyme activities with depth. Based on several studies, the rates (Davey et al. 2001; Zaccone et al. 2003) and spectra of enzyme activities (D’Ambrosio et al. 2014; Hoarfrost et al. 2017; Steen et al. 2012) decrease with depth, whereas cell-specific activities increase (Baltar et al. 2009). However, detection of a wider spectrum of polysaccharide-hydrolyzing enzymes at bottom waters in Guaymas Basin (Ziervogel and Arnosti, 2020) and at 500 m in the central Arctic (Balmonte et al. 2018) compared to those in surface waters demonstrate the nuances of these patterns. Greater similarity in enzyme activity depth profiles at a single station – than across locations – highlights surface-to-deep ocean connectivity and spatial trends even in the oceanic interior (Hoarfrost et al. 2017). With few enzyme activity measurements in deep waters, however, understanding of microbial control on organic matter degradation remains particularly limited in the mesopelagic and bathypelagic.

Delivery of organic matter from the surface to the deep ocean occurs in large part through the sinking of particulate organic matter (POM). During transport, POM can become fragmented through abiotic (i.e. shear stress) and biotic (e.g. enzymatic hydrolysis and zooplankton feeding) processes (Briggs et al. 2020; Zhao et al. 2020). Intense enzymatic hydrolysis of POM (Smith et al. 1992) – either via cell-bound or dissolved free enzymes – underscores the importance of particle-associated taxa for POM degradation (Baltar et al. 2010; Vetter et al. 1998, Zhao et al. 2020). A wider spectrum of enzyme activities has been detected on particles than either the free-living fraction (D’Ambrosio et al. 2014) or in bulk waters (Balmonte et al. 2018; Balmonte et al. 2020), observations that highlight the broad range of enzymes required to degrade POM. However, these investigations have only been carried out in a limited range of settings. Thus, possible variations in enzymatic capabilities of particle-associated taxa with latitude and depth, which would influence particle export and transfer efficiencies (Henson et al. 2012; Henson et al. 2019; Weber et al. 2016), remain underexplored.

Along a transect from 30°S in the South Pacific Gyre to 59°N in the Bering Sea, we investigated latitudinal and depth-related variations in the rates and substrate spectra of microbial enzyme activities in bulk seawater and on particles (≥ 3 μm). Based on previous latitudinal patterns of enzyme activities in surface waters (Arnosti et al. 2011), we hypothesized that the individual enzymes would exhibit varying patterns along latitudinal and depth gradients, likely due to varying sources, controls, and temperature optima. We additionally tested the hypothesis that patterns of particle-associated enzyme activities exhibit latitudinal and depth-related patterns that differ from those measured in bulk water (non-size fractionated), such that particle-associated taxa may be sources of distinct enzymatic activities. Using a suite of structurally-diverse substrates, we measured potential rates of frequently and infrequently-measured peptidases, glucosidases, and polysaccharide hydrolases. We identified differences in microbial capabilities to initiate organic matter degradation across water masses, and the extent to which differences in enzyme activities parallel emerging latitudinal patterns of microbial community structure and metabolic potentials.

## Materials and Methods

### Cruise track and sample collection

Samples were collected during the SO248 cruise (30°S to 60°N; Fig. 1a) on board R/V *Sonne* from May 3 to May 30, 2016. Bulk (non-size fractionated) water samples were collected from 19 stations at different depths using 20 L Niskin bottles mounted on a CTD rosette (Table S1). Additional water was collected to obtain particles via gravity filtration through 3 μm pore-size, 47 mm Millipore membrane filters; the volumes filtered for particle-associated analyses varied by depth and station (Table S2). All incubations (described below) and gravity filtration setups were kept either at room temperature (ca. 20°C), 15°C, or 4°C, depending on the ambient temperature of seawater (Table S1). Due to the substantial workload involved in the sample program, nine of the 19 stations were designated as ‘main stations’—locations at which peptidase, glucosidase, and polysaccharide hydrolase activities were measured at five depths. At all main stations, four depths were consistently sampled: surface, deep chlorophyll *a* maximum (DCM), 300 m, and 1000 m. For the three remaining main stations, the fifth depth was either at 500 m (S1 and S2) or at 75 m (S7) (Table S1). Additionally, particle-associated activities were also measured at all main stations at the DCM, 300 m, and 1000 m. At the 12 other stations, only peptidase and glucosidase activities were measured, and only in surface and DCM water.

**Figure 1.**
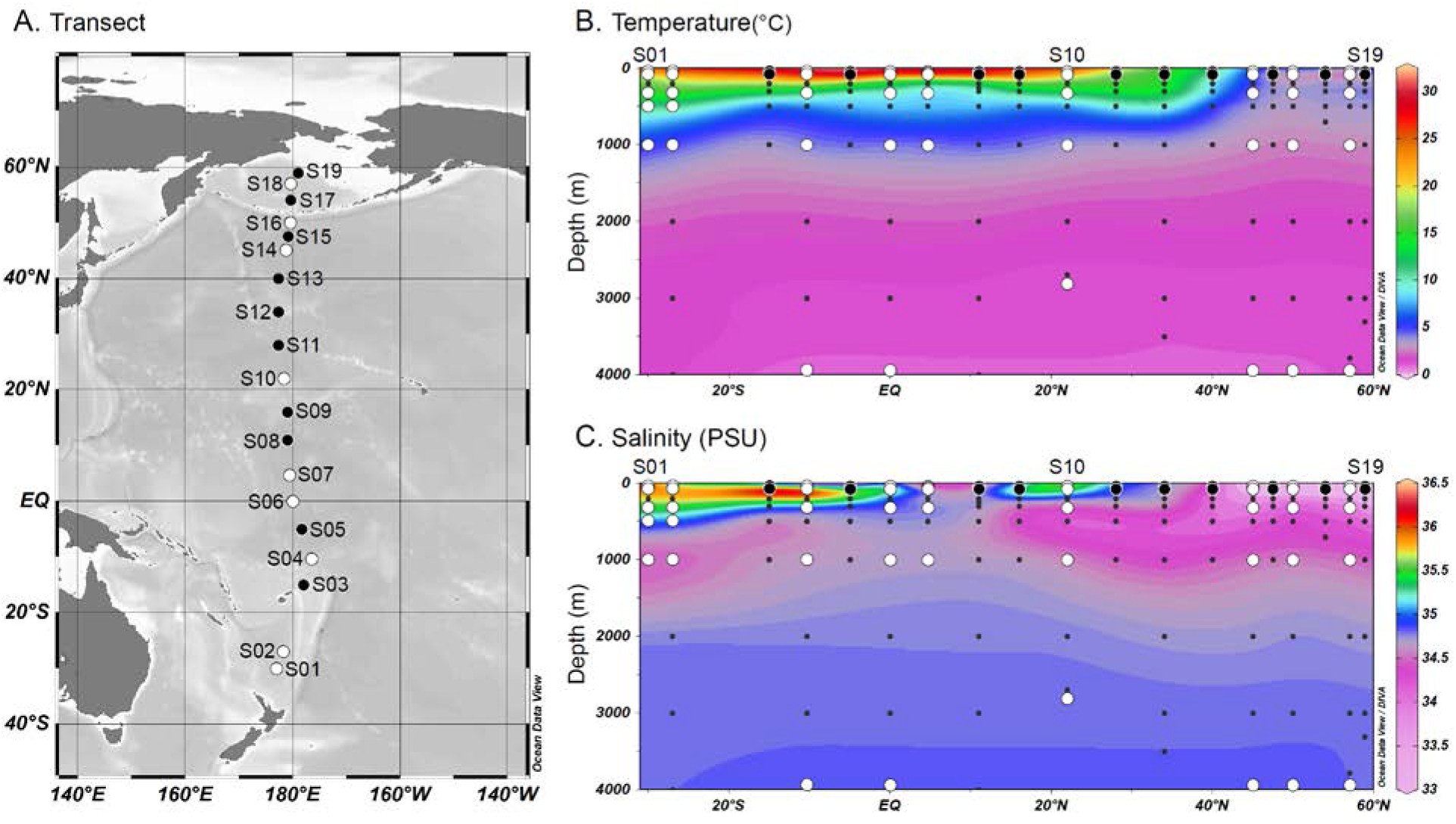
Map of stations from the South Pacific to the Bering Sea (A), temperature (B) and salinity (C) along the transect. Large black circles denote the stations where only bulk water peptidase and glucosidase were measured at the surface and DCM. White circles denote the stations where the full spectrum of enzymatic assays was carried out at all depths.

### Particulate organic carbon and nitrogen, chlorophyll a

Water samples (1.5 L to 6 L, depending on station and depth) were filtered onto pre-combusted (2 h, 450°C) and pre-weighed GF/F filters (Whatman) for particulate organic carbon (POC) and particulate organic nitrogen (PON) analysis. Filters were rinsed with distilled water to remove salt and kept frozen at −20°C until analysis as previously described (Lunau et al. 2006). To measure chlorophyll *a* (chl *a*), ca. 1 to 3 L of seawater was filtered through 25 mm GF/F filters (Whatman, Munich, Germany). After filtration, filters were wrapped in foil and stored at −80°C prior to analyses. Filters were crushed and extracted in 90% ice-cold acetone in the dark for 2 h. Concentrations were measured using a fluorometer (Turner Designs) and calculated according to established protocols (Tremblay et al. 2002). A standard solution of chl *a* was used for fluorescence calibration (Sigma-Aldrich).

### Bacterial abundance, production, growth rates, substrate turnover

Bacterial abundances were determined by SybrGreen I (Invitrogen) DNA staining on board using a BD Accuri C6 flow cytometer (Biosciences), after the protocol of Giebel et al. (2019). Bacterial biomass production rates were quantified using ^14^C-leucine incorporation (Simon and Azam, 1989; Simon et al. 2004). Briefly, 10 mL of triplicate water samples and a formaldehyde killed control (1% vol:vol) were incubated at *in situ* temperature with ^14^C-leucine (334 Ci mmol^−1^; Hartmann Analytics, Braunschweig, Germany) at a final concentration of 20 nM, and stored in the dark. After 2-10 h, formaldehyde was added to stop bacterial ^14^C-leucine incorporation. Samples were then filtered using 0.2 μm nitrocellulose filters (25 mm), extracted with ice cold 5% trichloroacetic acid, and analyzed by radioscintillation counting. Bacterial production rates were calculated using a conversion factor of 3.05 kg C mol^−1^ leucine^−1^ (Simon and Azam, 1989). Bacterial growth rates (μ day^−1^) were calculated as:

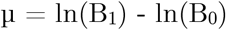

B_0_ and B_1_ (B_0_+BP) are bacterial biomass at t_0_ and t_1_, respectively. Bacterial biomass was calculated from bacterial cell numbers, assuming a carbon content of 20 × 10^−15^ g C cell^−1^ (Simon and Azam, 1989), and BP is bacterial biomass produced over 24 h and measured by leucine incorporation, as mentioned above.

Turnover rate constants of dissolved free amino acids (DFAA), acetate, and glucose were measured through the incorporation of a mix of ^3^H-DFAA (mean specific activity 2.22 TBq mmol-1, Hartmann Analytic), ^3^H-glucose (2.22TBq mmol-1, Hartmann Analytic) and ^3^H-acetate (0.925 TBq mmol-1229, Hartmann Analytic), following previously established procedures and calculations (Simon and Rosenstock, 2007). Note that these rates yield conservative values, as the assay captures only substrates incorporated into biomass and neglects that fractions taken up into the cytosol and respired.

### Bulk seawater enzyme activity assays

Peptide and glucose substrate analogs were used to measure peptidase and glucosidase activities, respectively (Hoppe, 1983). Exo-acting (terminal cleaving) leucine-aminopeptidase (leu-AMP) activities were measured using the methylcoumarin (MCA)-labelled substrate analog Leucine-MCA (Leu). Activities of the endo-acting (mid-chain cleaving) chymotrypsins (chym) and trypsins (tryp) were assayed using the following MCA-labeled substrates: Alanine-Alanine-Phenylalanine (AAF-chym), Alanine-Alanine-Proline-Phenylalanine (AAPF-chym), Phenylalanine-Serine-Arginine (FSR-tryp), and Butyloxycarbonyl-Glutamine-Alanine-Arginine (QAR-tryp). Glucosidase activities were measured using the following methylumbelliferyl (MUF)-labelled compounds: α-glucopyranoside (α-glu) and β-glucopyranoside (β-glu). Whereas Leu, α-glu, and β-glu have been used in a wide range of environmental settings, substrate proxies measuring chymotrypsin and trypsin activities have been used only in a limited number of systems (e.g., Arnosti, 2015; Balmonte et al. 2019; Balmonte et al. 2020; Bong et al. 2013; Obayashi and Suzuki, 2005; Steen and Arnosti, 2013).

Incubations for bulk water peptidase and glucosidase activity were set up in flat bottom, black 96-well plates. Triplicate wells were set up for live bulk water, as well as for killed controls prepared using cooled, autoclaved ambient seawater. Substrates – prepared in DMSO at a stock concentration of 5 mM – were added to the triplicate live and killed control wells (200 μL) to a final concentration of 100 μM, which we assumed was at or near substrate saturation based on previous studies using some of the same substrates in the subarctic Pacific (Fukuda et al. 2000). Fluorescence was measured using a Tecan Infinite F200 plate reader (Austria) with excitation and emission wavelengths of 360 and 460 nm, respectively, at several timepoints: immediately after substrate addition (t0), 12 h (t1), 24 h (t2), and 48 h (t3). Fluorescence values were converted to concentrations of hydrolyzed substrate using a standard curve of fluorescence vs. different concentrations of MCA or MUF fluorophores. Rates were normalized by the volume of incubation and averaged across triplicates. Only rates for t1 are reported in this study.

Fluorescently labeled polysaccharides were used to measure polysaccharide hydrolase activities (Arnosti 2003). These substrates include pullulan [α(1,6)-linked maltotriose], laminarin [β(1,3-glucose)], xylan (xylose), fucoidan (sulfated fucose), arabinogalactan (arabinose and galactose), and chondroitin sulfate (sulfated *N-*acetylgalactosamine and glucuronic acid) (Arnosti, 2003; Teske et al. 2011). The monomer constituents of these polysaccharides are widely detected in the marine water column (Benner, 2002), largely from algal and phytoplankton sources (Painter, 1983). Genes for enzymes that hydrolyze these polysaccharides have also been detected among various bacterial taxa (Alderkamp et al. 2007; Elifantz et al. 2008; Neumann et al. 2015; Teeling et al. 2012). Incubations to measure bulk water polysaccharide hydrolase activities follow an established protocol (Arnosti, 2003).

Triplicate incubations and a singleton killed control were prepared in 15 mL centrifuge tubes. The killed control was prepared using cooled, autoclaved ambient seawater. Polysaccharide substrates were added (one per tube) to a final concentration of 3.5 μM monomer equivalent, except for fucoidan, which was added to a final concentration of 5.0 μM monomer equivalent due to low labeling density of the polysaccharide. The incubations were subsampled at the following timepoints: immediately at substrate addition (t0), plus 5 d (t1), 10 d (t2), 15 d (t3), and 25 d (t4). To subsample the 15 mL incubations, 2 mL were collected from each incubation, and filtered through a 0.2 μm surfactant-free cellulose acetate (SFCA) filter. The filtrate was collected in a 2 mL centrifuge tube, and frozen at −20°C until further analysis. Changes in substrate molecular weight over time were measured via gel permeation chromatography, and hydrolysis rates were calculated as previously described (Arnosti, 2003). Only maximum potential rates, which occur at different timepoints, are reported in the main text; data for all timepoints are available as Supplementary Information (Fig. S6e-f).

### Particle-associated enzyme activity assays

Filters (3 μm pore size) used to collect particles via gravity filtration (see Section 2.1) were cut into evenly-sized 1/12^th^ pieces using sterile razor blades (Balmonte et al. 2018). Particle-associated enzyme activities (glucosidase, peptidase, polysaccharide hydrolase) were measured using two different incubation setups. Particle-associated peptidase and glucosidase activities measured in duplicate by submerging two particle-containing filter pieces (each 1/12^th^ of entire filter) in separate 4 mL cuvettes containing cooled, autoclaved ambient seawater. A single killed control was prepared by submerging a sterile filter piece (1/12^th^ of unused filter) in 4 mL of cooled, autoclaved ambient seawater. Substrates were added to a final concentration of 100 μM. At four timepoints – upon addition of substrate (t0), 24 h (t1), 48 (t2), and 72 h (t3) – live duplicates and killed control singleton were subsampled by pipetting 3 × 200 μL (for technical triplicates) per incubation from each 4 mL cuvette into a 96 well plate for fluorescence measurement. Fluorescence values were converted to hydrolysis rates as described above for bulk water enzymatic assays, but were normalized to the volume of filtrate that passed through the 3 μm filter (Table S2). Since hydrolysis was well advanced at t1, only values at 24 h (t1) were included in this study. The percentage of rates measured in bulk water that can be attributed to the particle-associated fraction was calculated using the following equation:

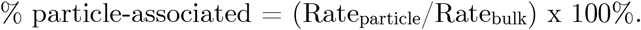

A value of 0 indicates that no rate for a specific enzyme was measured in the particle-associated fraction, whereas a value of 100% indicates that the rate measured for a specific enzyme was entirely particle-associated. A reported value of 100% may be due to three reasons: (1) enzyme-specific rates for bulk water and the particle-associated fraction were equivalent, (2) rates for bulk water were less than those for the particle-associated fraction, or (3) rates for bulk water were absent, and rates for specific enzymes were only detectable in the particle-associated fraction. While the resulting % particle-associated would be greater than 100% for scenario (2), we only report a maximum of F100%. For scenario (3), the equation would not be possible, in which case we manually inserted 100%.

Incubations to measure particle-associated polysaccharide hydrolase activities were set up in 15 mL centrifuge tubes filled with cooled, autoclaved ambient seawater (Balmonte et al. 2018). Filter pieces (1/12^th^) containing particles were submerged, and polysaccharide substrates were added to a final concentration of 3.5 μM monomer equivalent, with the exception of fucoidan (5 μM). Incubations were subsampled by drawing 2 mL, and filtering through a 0.2 μm SFCA filter. The filtrate was captured in a 2 mL centrifuge tube and stored at −20°C until further processing in-lab. As with the bulk rates, only the maximum potential particle-associated rates, measured at different timepoints, are discussed in this study; the remaining timepoint data are available in the Supplementary Information (Fig. S7f).

### Data visualization and statistical analyses

Ocean Data View (ODV) was used to create a station map, as well as to plot temperature and salinity. All enzyme activity and correlation plots were visualized on R using the package ‘ggplot2’ and ‘corrplot’, respectively. For bulk peptidase activities in surface waters and DCM, a curve was fitted, specifying the ‘loess’ model, and a 95% confidence interval was calculated. Non-metric multidimensional scaling (NMDS) through the R package ‘vegan’ was used to ordinate bulk versus particle-associated rates using the Bray-Curtis dissimilarity index; the statistical significance of the difference between these groups was tested using PerMANOVA (999 permutations) through the function ‘adonis’ in the R package ‘vegan’. To measure the β-diversity of peptidase and glucosidase activities per depth, the Bray-Curtis dissimilarities of sample data points (based on bulk or particle-associated enzymatic profiles) to the per-depth group mean centroid was calculated using the function ‘betadisper’ in the R package ‘vegan’. Correlation plots were based on the Pearson correlations between enzyme activities and multiple biotic and abiotic parameters. Correlation analyses were separately run using data from all depths, as well as data from individual depth realms (e.g. epipelagic, mesopelagic, and bathypelagic). Shannon indices were calculated based on an established equation fitted for enzyme activities (Steen et al. 2010).

### Data availability

Hydrographic data are available in the PANGAEA repository (Badewien et al. 2016). Enzyme activity data are available in the BCO-DMO database (Arnosti, 2020a, 2020b, 2020c, 2020d). Data for bacterial cell counts, bacterial production rates, growth rates, substrate turnover rates, POC and PON concentrations, and C:N ratios are available in the PANGAEA repository (Giebel et al. 2020). Only a portion of the dataset by Giebel et al. (2020)—from stations and depths where enzymatic activity data were measured (Table S1)—were used for this manuscript.

## Results

### Environmental context

The transect stations covered a wide range of environmental gradients (Fig. 1a). Surface water temperatures varied from 4.2°C at S18 (57°N) to 30.4°C at S05 (5°S) (Fig.1b). Surface water salinity ranged from 32.90 PSU at S19 (58.9°N), up to 35.92 PSU at S01 (30°S) (Fig. 1c); Table S1). At 300 m and below, temperature was highest (16.1°C) at 300 m at S02 (27°S) and lowest (1.0°C) at 4000 m at S04 (10°S). POC in surface waters covered a narrow range of ca. 28-32 μg L^−1^ from 20°S to 20°N (Fig. S1a). POC increased gradually in the North Pacific Subtropical Gyre, peaking at ca. 160 μg L^−1^ at the southern edge of the North Pacific Polar Frontal zone (NPPF; 40°N), comparable to the highest value at 60°N in the Bering Sea. POC at the DCM paralleled surface water trends from 20°S to 20°N, but values did not peak with the same intensity at 40°N (Fig. S1a). POC at the DCM reached its highest concentration also at 59°N, but was only ca. 103 μg L^−1^. In the surface and DCM, PON trends mirrored surface water POC trends (Fig. S1b), yielding relatively consistent POC:PON ratios in surface waters and at the DCM (Fig. S1c). In mesopelagic waters (300 m) and below, POC and PON concentrations were low (Fig. S1a,b), but with POC:PON ratios higher than in the epipelagic (Fig. S1c).

### Bacterial counts, production, and growth

In surface waters, total bacterial cell counts were highest in the northernmost latitudes, reaching values of 1.81 × 10^6^ cells mL^−1^ at 40°N, and peaking at 2.17 × 10^6^ cells mL^−1^ at 54°N in the Bering Sea. At the DCM, bacterial cell counts were comparable to those in surface waters from 30°S until 40°N, but became decoupled further north. Highest bacterial cell counts at the DCM were measured at 40°N, at ca. 1.04 × 10^6^ cells mL^−1^. At depths of 300 m and below, cell counts were generally an order of magnitude lower than values from the epipelagic (Fig. S1d).

Bacterial production patterns deviated from bacterial cell count trends. In surface waters, two peaks were observed for bacterial production: ca. 1170 ng C L^−1^ hr^−1^ at 5°S (S05), and ca. 1282 ng C L^−1^ hr^−1^ at 16°N (S09) (Fig. S1e). At the South Pacific Subtropical Gyre (30°S to 10°S), bacterial production remained low, ranging from 0.3 to 15.7 ng C L^−1^ hr^−1^. Bacterial production rates north of the second peak were in the range of 50.3 to 321 ng C L^−1^ hr^−1^. Bacterial production rates at the DCM throughout the transect remained low to moderate, only reaching ca. 248 ng C L^−1^ hr^−1^. Trends for bacterial growth rates paralleled those for bacterial protein production (Fig. S1f).

### Latitudinal trends in bulk peptidase and glucosidase activities

Peptidase and glucosidase activities showed distinct latitudinal trends at the surface and DCM (Fig. 2a). Leu-AMP exhibited strong latitudinal variation, with lowest activities in the North Pacific subtropical gyre (15°N), and gradually increasing activities with increasing latitude (Fig. 2a), which peaked in the Bering Sea (Fig. S2a). The positive correlation of leu-AMP activities with latitude was higher at the surface (R^2^ = 0.77, p < 0.001) than at the DCM (R^2^ = 0.46, p < 0.001). An even stronger — but negative — correlation was observed between leu-AMP activities and temperature, both for surface (R^2^ = 0.83, p < 0.001) and DCM (R^2^ = 0.49, p < 0.001). No other peptidase or glucosidase activities exhibited a significant relationship with latitude. Moreover, leu-AMP (exopeptidase) activities exhibited gradual transitions with latitude, whereas the chymotrypsin and trypsin (endopeptidase) activities were highly patchy. Substantial patchiness is evident in the broad 95% confidence interval for the fitted curve for the endopeptidases (Fig. 2a). All endopeptidase activities exhibited distinct latitudinal patterns, even those within the same enzyme class (e.g. AAF-chym vs. AAPF-chym, QAR-tryp vs. FSR-tryp), and display decoupled trends between surface waters and the DCM (Fig. 2a).

**Figure 2.**
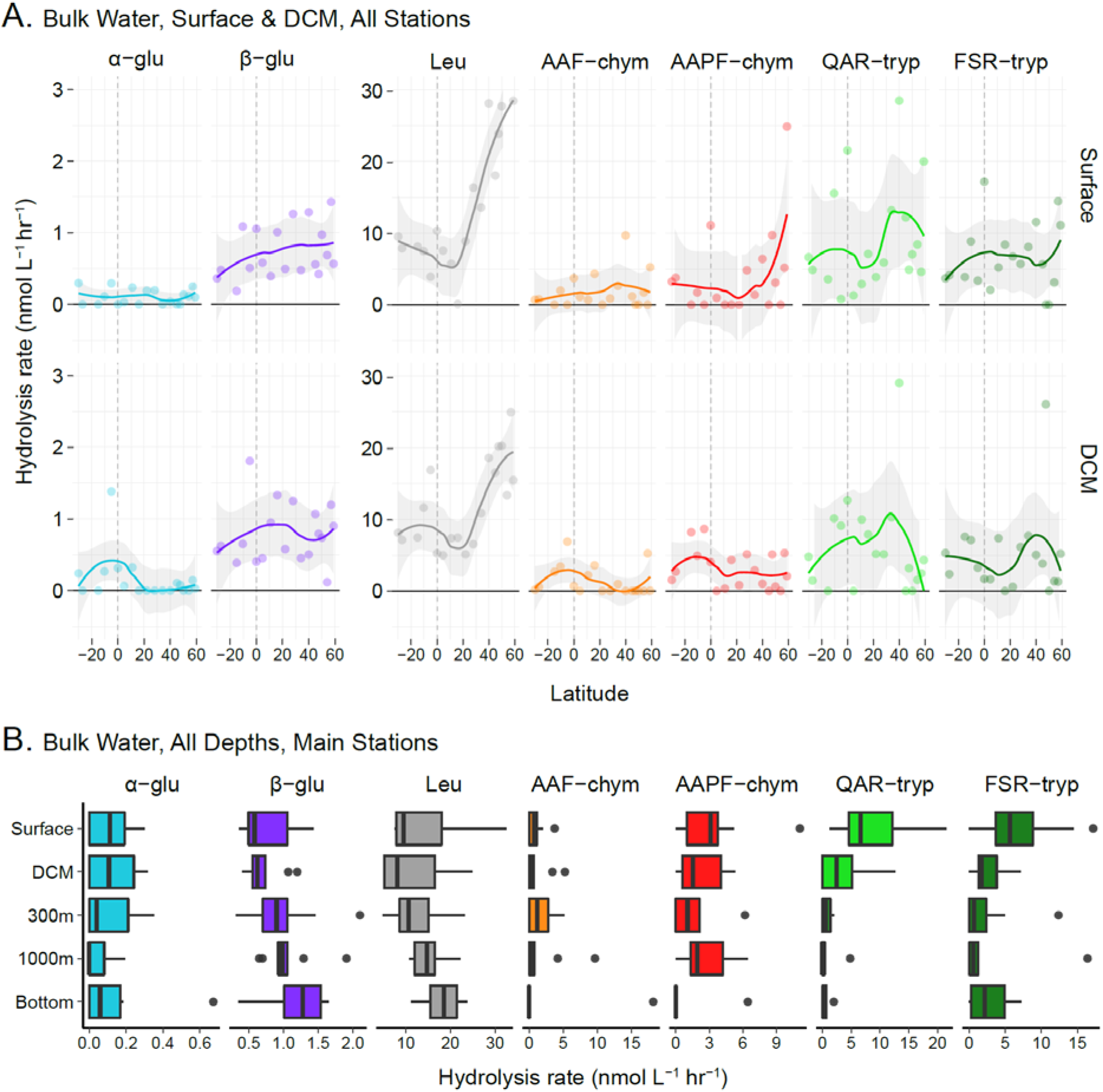
Bulk (non-size fractionated) glucosidase and peptidase activities in surface waters and at the deep chlorophyll maximum (DCM) at all stations (A), and at all depths for the main stations (B). Negative latitudes are those south of the equator. Note that DCM and bottom water depths vary by station, and x-axis scales differ by substrate. Gray shading (A) represents 95% confidence interval.

Endopeptidase patchiness largely drives differences in enzymatic spectra (Fig. S2a) and summed (combined) peptidase and glucosidase activities (Fig. S2b). For example, in the equatorial surface water (S06), all endopeptidases showed moderate to high activities, but this station was adjacent to two stations in which all endopeptidases either exhibited low or no activities (Fig. S2a). Hence, patchiness among summed rates is also observed, with the three highest values in surface waters observed at the northernmost station in the Bering Sea (60°N, S19), 40°N (S13), and at the equator (Fig. S2b).

### Depth-related trends in bulk peptidase and glucosidase activities

Substantial variations in enzymatic profiles were apparent with increasing depth. Within the epipelagic, surface and DCM patterns were frequently decoupled (Fig. 2a, S2a, S2b). For instance, low rates at 5°S (S05) surface water contrasted with the high rates of a wide range of peptidases and glucosidases at the DCM (Fig. S2a). Moreover, consistently high summed rates in surface waters and DCM were only observed at 40°N (S13); the two other peak summed rates at the DCM were measured at 47.5°N (S15) and 5°S (Fig. S2b).

The spectrum of enzyme activities became more limited with increasing depth (Fig. S3a), and latitudinal patterns observed in the upper water column were also attenuated at depth (Fig. S3b). With few exceptions (Fig. S3b), this trend resulted in generally lower summed rates (Fig. S3c) and Shannon indices (Fig. S3d) in the deepest waters, driven most proximately by decreasing rates of endopeptidase activities (Fig. 2b), often to undetectable levels (Fig. S3a). In contrast, exo-acting leu-AMP and β-glucosidase activities in bottom waters were measured at rates either comparable to — or higher than — those in the upper water column (Fig. 2b, S3a). Hence, depth-averaged rates of glucosidase and peptidase activities demonstrate notable enzyme-specific patterns with increasing depth (Fig. 2b).

### Particle-associated versus bulk water peptidase and glucosidase trends

Enzyme activities on particles were distinct from those detected in bulk waters (Fig. 3a) (PerMANOVA, Bray-Curtis, R^2^ = 0.37, p < 0.001). Bulk water and particle-associated enzyme activities also became more dissimilar with increasing depth, evident in the ordination as overlapping data points at the DCM, but near-complete separation of points at 1000 m (Fig. 3a). Particle-associated enzyme activities exhibited more variability—visible with loose clustering of data points—than patterns observed in bulk water (Fig. 3a). This activity variability on particles increased deeper in the water column, but bulk water enzyme activities showed the opposite trend (Fig. 3b). Results were similar when comparing bulk water and particle-associated results both at 24 h (Fig. 3a-c), or at 12 h and 24 h, respectively (Fig. S4a-c).

**Figure 3.**
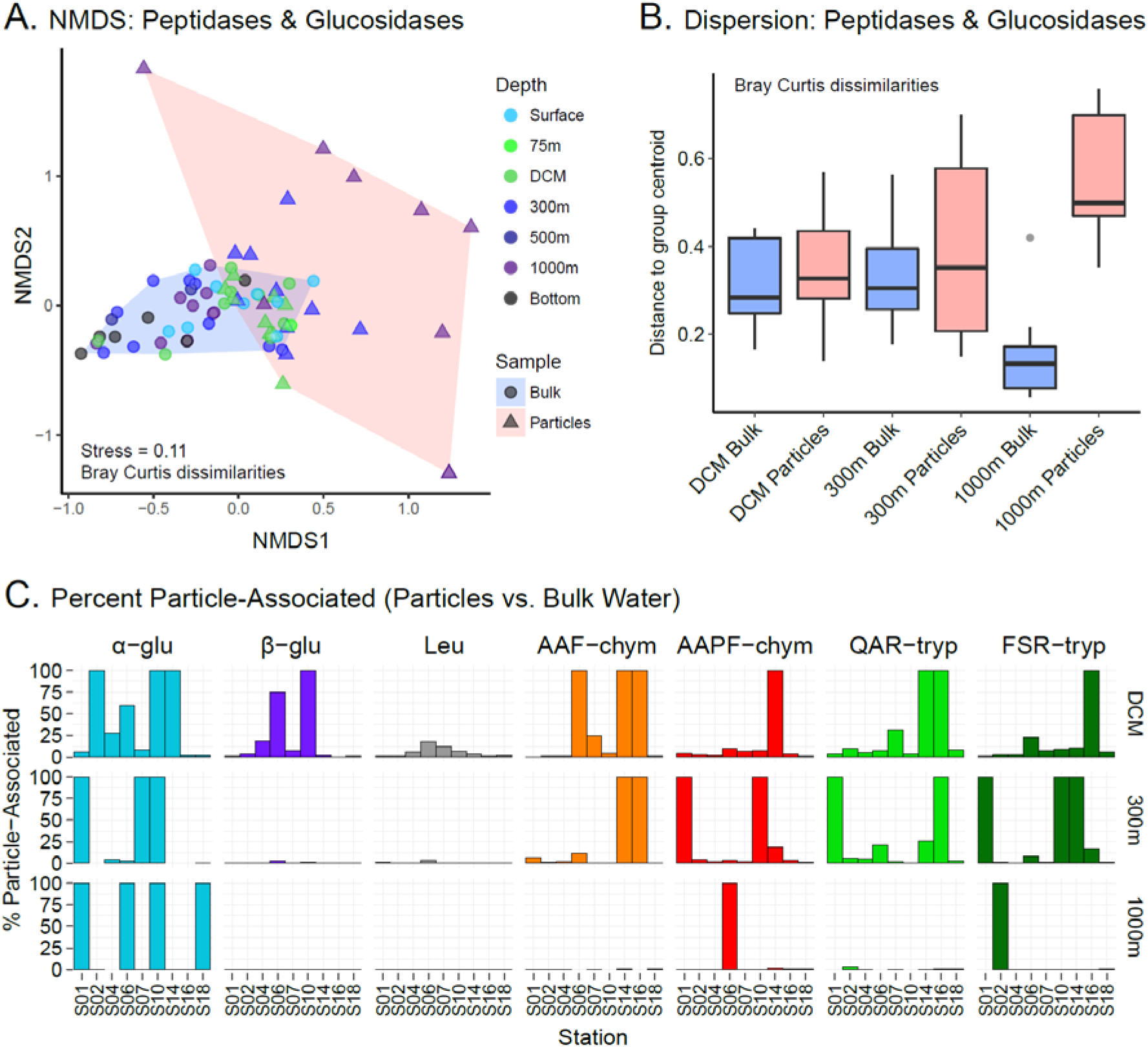
Non-metric multidimensional scaling (NMDS) plot of bulk water (blue) and particle-associated (pink) glucosidase and peptidase activities using the Bray-Curtis dissimilarity index (A). Bray-Curtis based group dispersions of bulk water and particle-associated glucosidase and peptidase activities, measured as distances of sample rates to group centroid at the DCM, 300 m (mesopelagic), and 1000 m (bathypelagic) (B). Proportion of bulk water glucosidase and peptidase activities attributable to the ≥ 3 μm particle-associated fraction (C). Note that bulk water peptidase and glucosidase rates used here were measured at the 24 h (t2) timepoint, for direct comparison to the particle-associated rates at the 24 h (t1) timepoint.

Quantifying the percent contribution of particle-associated hydrolysis rates to bulk hydrolysis rates revealed substantial differences in relative proportions of most enzyme activities on particles compared to the bulk water. Whereas leu-AMP, at most, was ca. 18% particle-associated at the equator, glucosidase and endopeptidase activities were 100% particle-associated in some locations and depths (Fig. 3c). Remarkably, the most frequently-detected enzyme activities on particles at 1000 m were α-glucosidase, AAPF-chym, FSR-tryp, albeit at very low rates (Fig. S5a). The high relative importance of α-glucosidase and endopeptidases on particles persisted throughout the entire latitudinal transect.

Most particle-associated enzyme activities peaked in warm tropical and sub-tropical waters (Fig. S5a) and decreased with increasing latitude, mirroring trends in bulk water. High rates and a wider spectrum of enzyme activities were observed at the equator (S06) and at 5°N (S07) (Fig. S5b). Summed particle-associated activities in the DCM and at 300 m were highest at the equator and decreased towards the poles (Fig. S5c). Summed rates also declined with depth, accompanied by lower Shannon values (Fig. S5d); however, the steepest depth-related decreases were observed in equatorial waters. At 1000 m, peptidase and glucosidase activities were patchy, with no easily distinguishable spatial trend (Fig. S5a,b).

### Latitudinal variations in bulk polysaccharide hydrolase activities

Robust differences in polysaccharide hydrolase activities were observed throughout the transect, with the most prominent shift at 45°N (S14), within the NPPF (Fig. 4). At this station and those further north, enzymatic profiles were marked by higher chondroitin sulfate hydrolase activities than observed elsewhere (Fig. 4, S6a-b). As a consequence, chondroitin hydrolysis rates correlated positively with latitude (R^2^ = 0.43, p < 0.001). In contrast, laminarinase activities peaked in equatorial and adjacent waters (S01-S10), decreased towards the northernmost latitudes, and correlated positively with temperature at all depths (see Section 3.9). Xylan was hydrolyzed most rapidly in the low and mid-latitudes, but exhibited no significant relationship with latitude. The highest summed polysaccharide hydrolase rates in surface waters were observed at the equator (S06), due in large part to high laminarinase and xylanase rates (Fig. S6b). Pullulanase activities were measurable at most stations down to depths of 300 m, but also did not feature a strong latitudinal trend (Fig. S6a). Neither arabinogalactan nor fucoidan were hydrolyzed in bulk waters. Thus, polysaccharide hydrolases demonstrate individual latitudinal patterns, similar to findings for peptidases and glucosidases.

**Figure 4.**
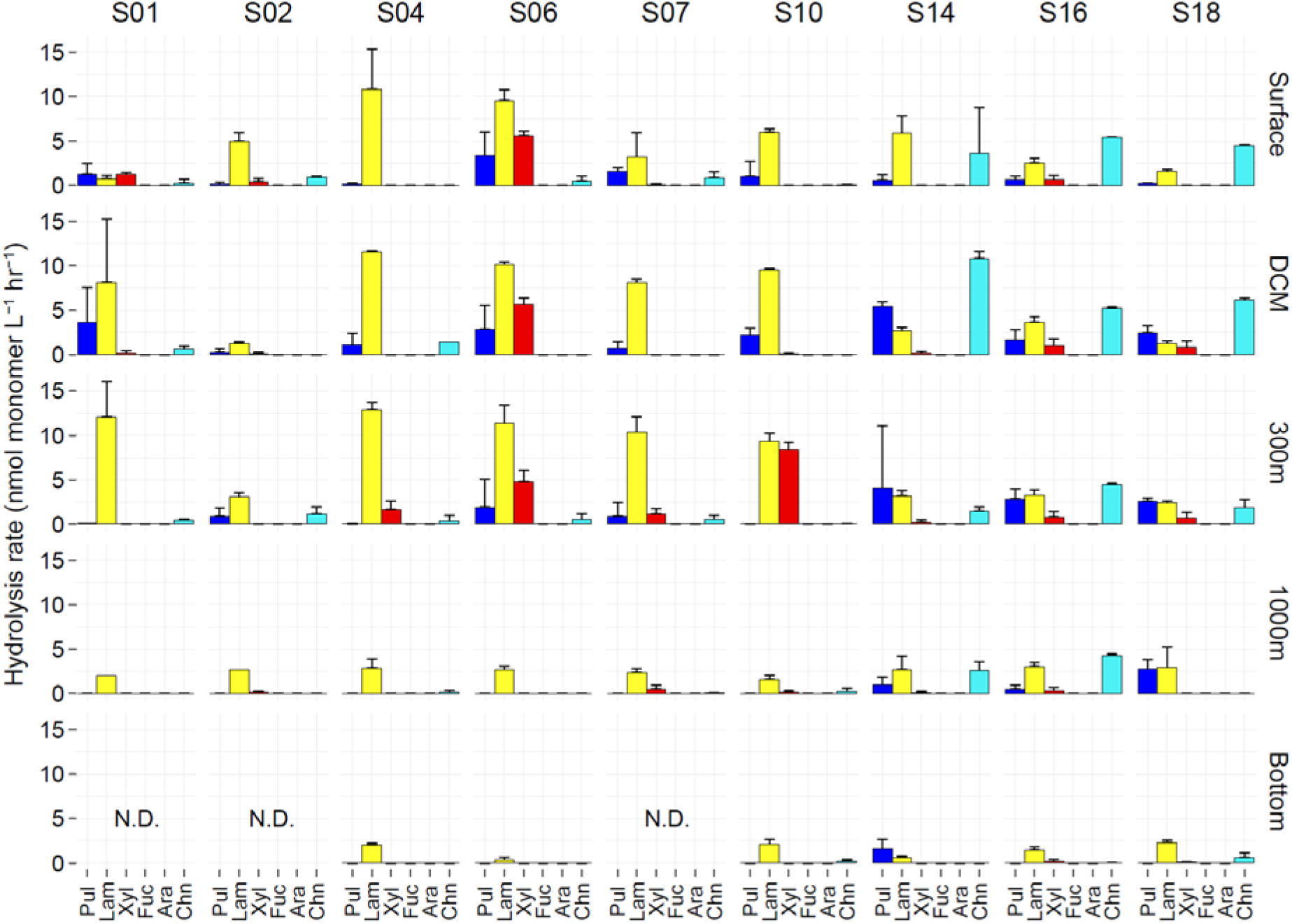
Polysaccharide hydrolase activities in bulk water. Note that data presented are maximum rates, which occurred at different timepoints. Pul = Pullulan, Lam = Laminarin, Xyl = Xylan, Fuc = Fucoidan, Ara = Arabinogalactan, Chn = Chondroitin Sulfate, N.D. = No Data. Error bars represent standard deviation between replicates.

### Depth-related differences in bulk polysaccharide hydrolase activities

Lower rates and more limited spectra of polysaccharide hydrolases characterized deep versus surface waters (Fig. 4, S6b). Summed rates remained comparable from surface waters to 300 m — at times with the highest summed rates detected at the DCM or at 300 m (Fig. S6b). Accordingly, highest Shannon values were frequently observed at the DCM or 300 m (Fig. S6d). Moreover, the depth of steep attenuation of rates and enzymatic spectra varied by latitude. In the low to mid-latitudes (S01-S10), rates and enzymatic spectra decreased markedly from 300 m to 1000 m, whereas those in high latitudes remained comparable over the same depth range (Fig. 4, S5b).

### Particle-associated polysaccharide hydrolase activities

The relative contributions of polysaccharide hydrolases differed on particles (Fig. 5) compared to bulk waters (PerMANOVA, Bray-Curtis, R^2^ = 0.39, p < 0.001), despite not being visible in the NMDS ordination (Fig. S7a,b). Wider spectra of polysaccharide hydrolases were measured on particles at the same station and depth (Fig. 5). This trend was largely concentrated in low latitudes and especially pronounced at 300 m at S02, and at the DCM and 300 m at S07—locations in which fucoidan was hydrolyzed on particles but not in bulk water (Fig. 5). Moreover, at the DCM at S07, five polysaccharides were hydrolyzed on particles, whereas only two were hydrolyzed in bulk water (Fig. 4). Less pronounced examples, which nevertheless demonstrate wider spectra on particles, were evident throughout the low latitudes.

**Figure 5.**
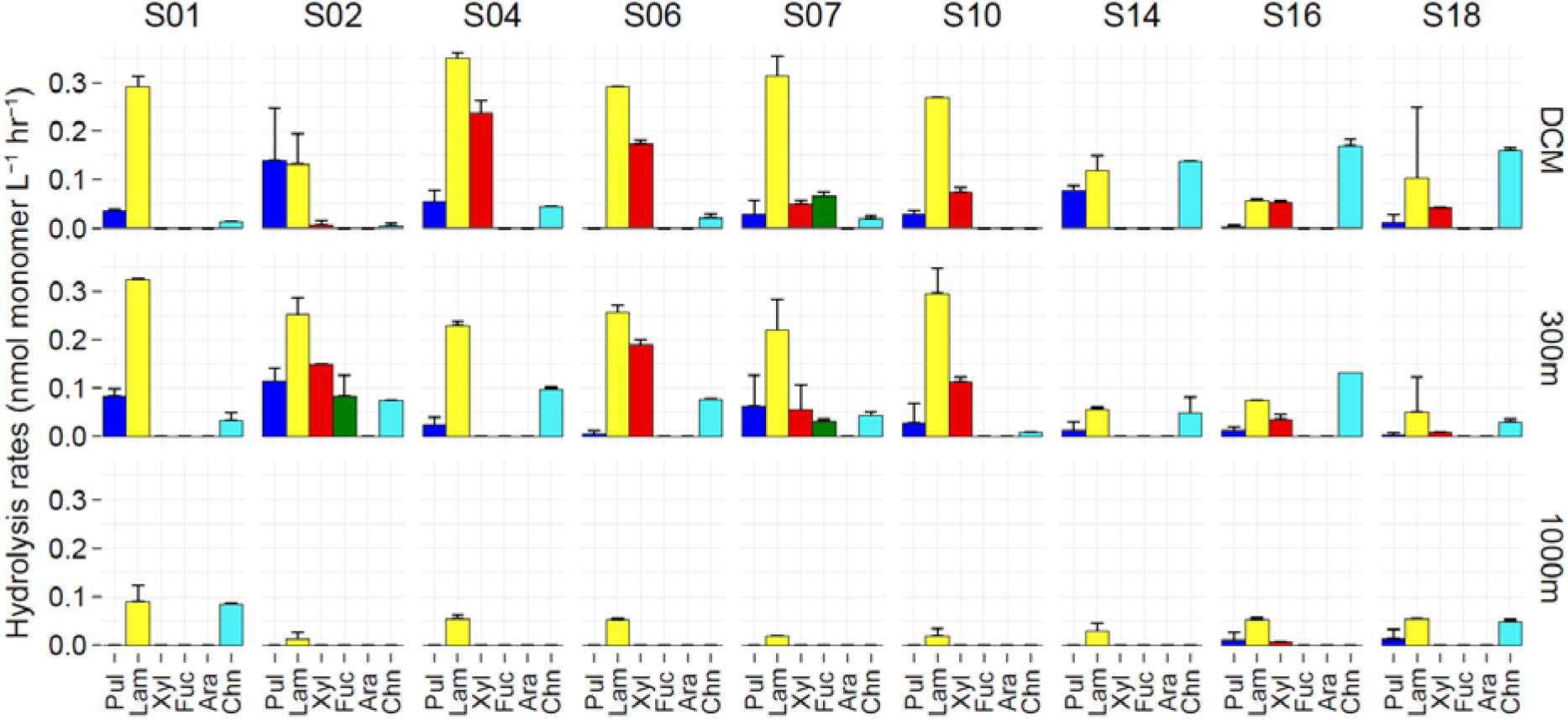
Polysaccharide hydrolase activities on ≥ 3μm particles. Note that data presented are maximum rates measured at different timepoints. Rates were normalized by volume of water filtered to obtain particles. Pul = Pullulan, Lam = Laminarin, Xyl = Xylan, Fuc = Fucoidan, Ara = Arabinogalactan, Chn = Chondroitin Sulfate. Error bars represent standard deviation between replicates.

Latitudinal and depth-related variations were observed for particle-associated activities, based on entire spectra (Fig. 5, S7c) and for individual enzymes (Fig. S7d). Summed rates in the DCM were higher in low latitudes than in high latitudes, with a peak at 10°S (S04) (Fig. S7c). With increasing depth, summed rates decreased, although at several stations these values were higher at 300 m than in the DCM (Fig. S7c). Enzyme-specific latitudinal trends were also observed on particles, and featured higher laminarinase and xylanase activities in low latitudes, but high chondroitin activities in high latitudes, particularly in the DCM (Fig. S7d). At 1000 m, only laminarinase activities were consistently detected; activities of other polysaccharide hydrolases were rarely detected. This increasingly limited spectrum of particle-associated activities (Fig. 5) parallels the depth-related trend observed in bulk water, for polysaccharide hydrolases, as well as peptidases and glucosidases.

As with patterns observed for particle-associated peptidases and glucosidases, the depth of steep attenuation of rates and enzymatic spectra for particle-associated polysaccharide hydrolases show pronounced latitudinal differences. High rates and wide spectra of particle-associated polysaccharide hydrolases in low latitudes—particularly visible from 27°S to 22°N (S02-S10)—were sharply attenuated from 300 m to 1000 m (Fig. 5). In contrast, particle-associated rates in the DCM and at 300 m in the high latitudes were comparable to those at 1000 m, and with little loss of polysaccharide hydrolase activities, particularly in the two northernmost stations (S16 and S18) (Fig. 5).

### Abiotic and biotic correlates of enzyme activities

Correlations between enzyme activities and physicochemical and bacterial parameters demonstrated varying trends based on enzyme class (e.g., peptidases, glucosidases, and polysaccharide hydrolases), activity source (i.e. bulk water vs. particles) and spatial scale (i.e., all depths vs. individual depths). Using data from all depths, more statistically significant abiotic and biotic correlates were identified for peptidases and glucosidases (Fig. 6a, 6b) than for polysaccharide hydrolases (Fig. 6c, 6d). Leu-AMP in bulk waters exhibited more significant correlations than other peptidases and glucosidases (Fig. 6a). Among the polysaccharide hydrolases, chondroitin sulfate hydrolysis correlated with the most variables (Fig. 6c). Particle-associated peptidases and glucosidases exhibited positive relationships with temperature, fluorescence, cell counts, and bacterial production (Fig. 6b), as well as co-occurrence patterns, evident by positive relationships with each other; such a trend was not observed among particle-associated polysaccharide hydrolases. Finally, most enzyme activities were decoupled from turnover of amino acids, glucose, and acetate (Fig. 6a-d).

**Figure 6.**
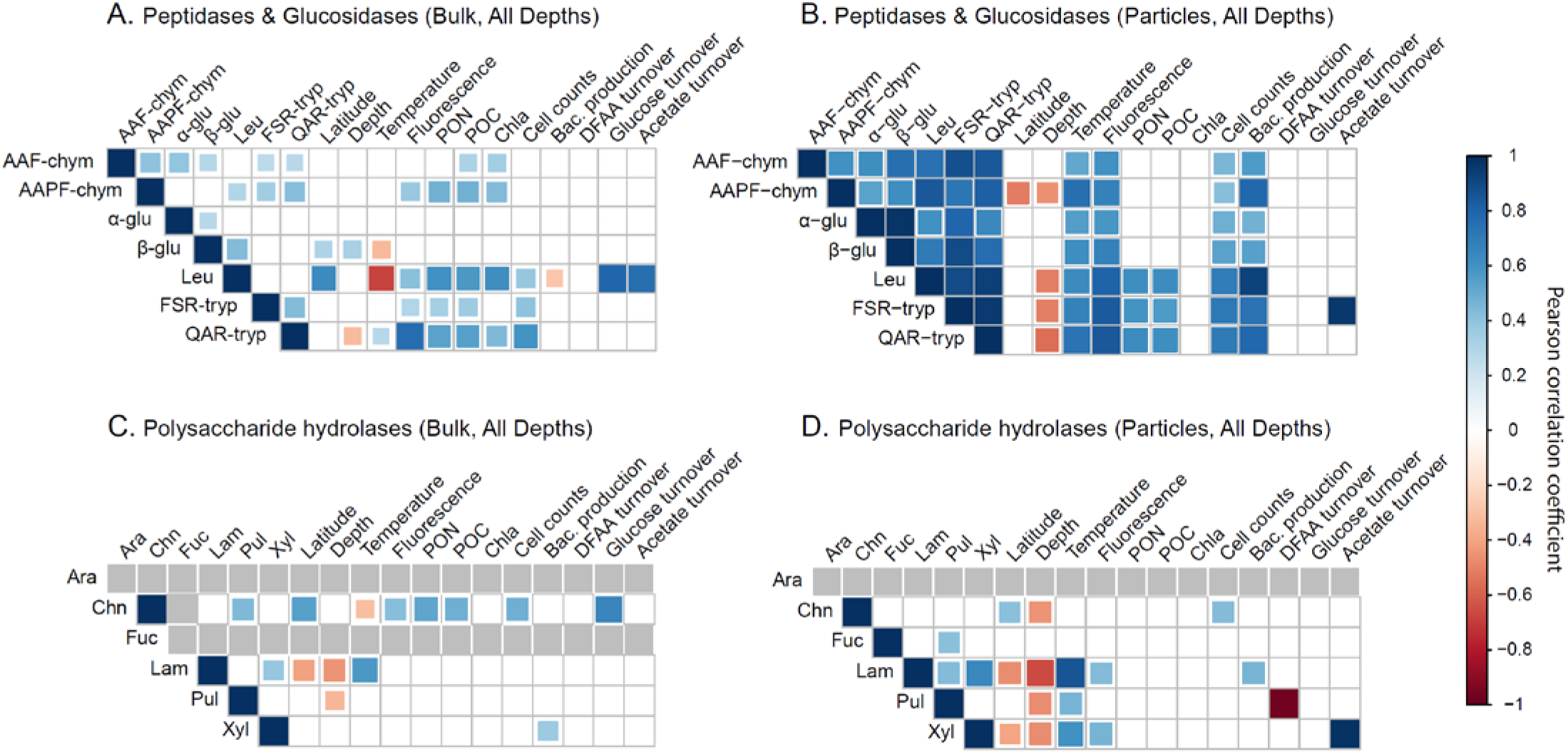
Pearson correlations between enzyme activities measured environmental and bulk bacterial activity parameters at all depths, separated by enzyme class and sample type: bulk water (A) and particle-associated (B) glucosidases and peptidases, as well as bulk water (C) and particle-associated (D) polysaccharide hydrolases. Non-significant correlations (p < 0.05) are shown as empty boxes. No rate data due to undetectable enzymatic activities are shown as gray boxes. Rates analyzed for panel (A) include bulk water rates from all stations.

As a caveat, many of the correlations identified using data from all depths (Fig. 6a-d) persisted or differed from those observed when data were sub-divided by different depths (i.e. epipelagic, mesopelagic, bathypelagic) (Fig. S8-S11), illustrating the scale dependence of these relationships. Numerous correlations of leu-AMP with biotic and abiotic parameters persisted throughout the water column, the most consistent of which is its negative relationship with temperature (Fig. S8a-d). Many of the correlates for particle-associated peptidases and glucosidases, as well the positive co-occurrence patterns, identified at all depths were undetected in the depth-separated analyses (Fig. S9a-d). Relationships between latitude and rates of several particle-associated peptidases and glucosidases were negative in datasets from all depths (Fig. 6b) and separately from the epipelagic (Fig. S9b) and mesopelagic (Fig. S9c), but positive in the bathypelagic (Fig. S9d). Among polysaccharide hydrolases, both in bulk waters (Fig. S10a-d) and on particles (Fig. S11a-d), few parameters correlated with enzyme activities; the range and strength of correlates even decreased with increasing depth. However, the most robust correlation was the consistent positive relationship of laminarinase activities with temperature, observed in all iterations of the analysis (Fig. S10a-d, S11a-d).

## Discussion

### Heterogeneous latitudinal and depth-related trends

Marine microbial communities exhibit substantial latitudinal and depth-related heterogeneity in their enzymatic capabilities to initiate OM degradation across a 9,800 km transect between the South Pacific Gyre and the Bering Sea. This “sea change” in enzymes across latitudes, depth, and the distinctions in bulk water vs. particle-associated patterns is summarized in Figure 7. Latitudinal trends are enzyme-specific (Figs. 2, 4): With increasing latitude, chondroitinase activities increase, but leucine aminopeptidase and laminarinase activities decrease (Fig. 7); activities of other peptidase and polysaccharide hydrolase exhibit significant spatial patchiness (Fig. 2-5). Enzymatic spectra — that is, the range of measurable activities at a given location (e.g., Fig. S2a, 4, 5) – become narrower with increasing latitude and depth (Fig. 7). These latitudinal (Arnosti et al. 2011) and depth-related trends (Hoarfrost et al. 2017; Balmonte et al. 2018), previously observed in bulk water, are also demonstrable in the particle-associated fraction. More importantly, differences in enzymatic activities in bulk water and on particles increase with increasing depth (Fig. 3b, 7), indicating that particle-associated taxa produce a set of enzymes that differ in range and proportions from those of their free-living counterparts. This multidimensional view of spatial heterogeneity in enzymatic patterns (Fig. 7) suggests different sources, controls, and substrate specificity of microbially-produced enzymes across considerable depths and distances.

**Figure 7.**
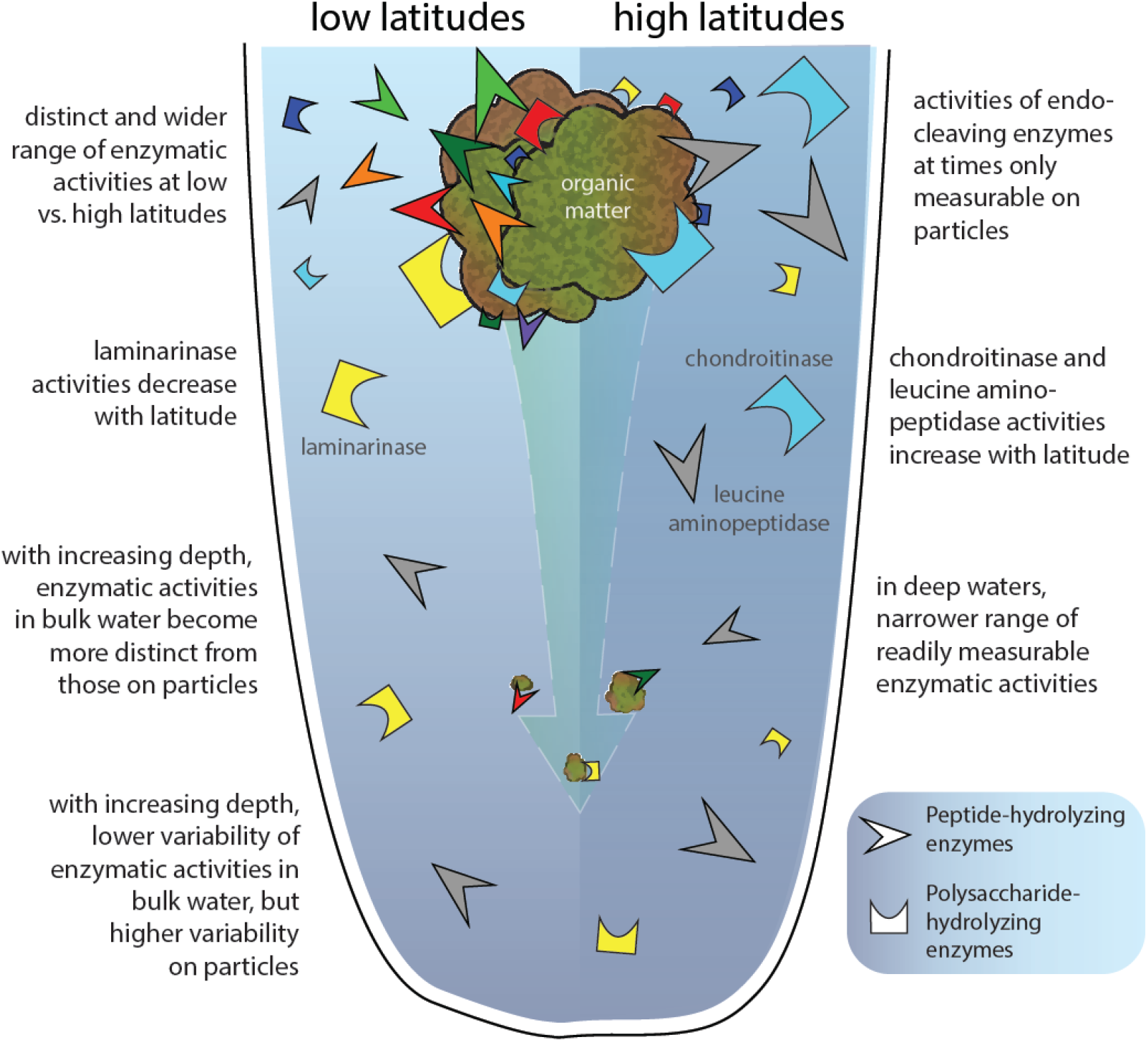
A Sea Change in Microbial Enzymes – a summary of trends in microbial enzymatic activities in bulk water and on particles in the open ocean, which differ in spectra and relative importance along latitudinal and depth gradients. Enzyme classes (i.e. peptidases vs. polysaccharide hydrolases) are represented by different shapes, and colors are consistent with the substrates used to measure their activities (Figs. 2-5).

Biotic and abiotic correlates for enzyme activities provide hints of potential controls and sources that differ both within and across enzyme classes, and on particles versus in bulk water (Fig. 6). Peptidase and glucosidase activities more frequently exhibit significant correlations with measured physicochemical and bacterial parameters (Fig. 6a,b) than do polysaccharide hydrolase activities (Fig. 6c,d). When all depths are considered, several peptidase and glucosidase activities correlate with factors related to primary production, including particulate organic carbon, particulate organic nitrogen, and chl *a* concentrations (Fig. 6a,b). Particle-associated peptidase and glucosidase activities across all depths (Fig. 6b) positively correlate with cell counts and bacterial production. These findings suggest that particulate matter derived largely from primary production is broken down to dissolved substrates (Fig. 7) and fuel biomass production of free-living bacteria that dominate in particle-poor pelagic environments (Smith et al. 1992). Moreover, variations in enzyme activities thus cannot be explained by individual variables, or by a common set of parameters. This result is consistent with previous findings from a large latitudinal gradient in surface waters (Arnosti et al. 2011) and in depth transects along a shorter latitudinal gradient in the Atlantic (Hoarfrost and Arnosti, 2017).

Correlates are rarely observed for polysaccharide hydrolases, suggesting that other factors likely better account for differences in these enzyme activities. The ability to degrade and utilize polysaccharides is a complex trait, requiring more genes to carry out these functions than simpler traits (Berlemont and Martiny, 2016). As a consequence, distribution of polysaccharide utilization among microorganisms is phylogenetically narrow – or restricted to a limited range of taxa. Hence, differences in microbial community composition may correspond to variations in polysaccharide hydrolase activities measured in low vs. high latitudes (Fig. 7). Latitudinal differences in marine microbial communities, both in structure and in function (Ghiglione et al. 2012; Sul et al. 2013; Ibarbalz et al. 2019; Salazar et al. 2019) support this explanation. In particular, rapid changes in epipelagic microbial community metagenomes and metatranscriptomes were detected at 40°N (Salazar et al. 2019) in the transition from thermally-stratified to well-mixed polar waters (Behrenfeld et al. 2006). Similarly, polysaccharide hydrolase profiles shifted markedly along the transect between the subtropical/temperate North Pacific and the sub-Arctic Pacific (S14, 45°N; Table S1), characterized by high chondroitin sulfate hydrolase and low laminarinase activities (Figs. 4, 5, 7). While the ecological boundary was identified among stations predominantly in the North Atlantic (Salazar et al. 2019), the transition from thermally-stratified to a well-mixed regime is also evident in the Pacific Ocean temperature profiles (Fig. 1b). Thus, a similar ecological boundary for microbial communities in the Pacific Ocean may have influenced spatial patterns of polysaccharide hydrolase activities.

Notably, the rates and spectrum of peptidase, glucosidase and polysaccharide hydrolase activities (Fig. 4, 5, S3a, S5b) decrease with increasing depth, and individual enzymes exhibit distinct patterns (Fig. 2b). Lower rates and limited enzymatic spectra in deeper waters (Fig. 7) are in accordance with previous enzyme activity depth profiles measured in the South and Equatorial Atlantic Ocean (Hoarfrost et al. 2017), Gulf of Mexico (Steen et al. 2012), and the central Arctic (Balmonte et al. 2018). These depth-related enzymatic patterns may indicate a widespread microbial strategy: compared to their deep water counterparts, microbial communities in surface waters invest greater resources to readily produce a more diverse set of enzymes to access the frequently-replenished organic matter supply in the upper water column (Fig. 7). However, microbial potential to produce many of these enzymes (Balmonte et al. 2019) and their substrate transporters (Bergauer et al. 2018; Zhao et al. 2020) exist in deep waters as in surface waters. The extent to which these enzymes are produced is therefore partially controlled by the availability of organic matter exported from surface waters (Fig. 7).

### Contrasting enzyme patterns on particles

Distinct proportions and often wider enzymatic spectra are detected on particles (Fig. 4, S5b) than in bulk seawater (Fig. 5, S3a), indicating the importance of particle-associate taxa for enzyme production even down to the bathypelagic (Fig. 7). Detection of genes encoding secretory CAZymes and peptidases, as well as transcripts and proteins for OM hydrolysis that likely belong to particle-associated taxa in the bathypelagic support this interpretation (Zhao et al. 2020). While the contribution of cell-bound vs. dissolved secreted enzymes cannot be ascertained from our measurements, a previous study suggests that much of the particle-associated enzymatic activities are likely due to enzymes secreted into the particle matrix (Zhao et al. 2020). Further, endo-acting peptidase activities can at times be attributed only to the particle-associated fraction (Fig. 3c, S5b). In contrast, only a minor fraction of exo-acting peptidase activities—measured here using the leucine substrate—is detected on particles (Fig. 3c). Endo-acting enzymes thus likely play a critical role in degradation of particles, as previously observed among polysaccharide hydrolases in the North Atlantic (D’Ambrosio et al. 2014), the central Arctic (Balmonte et al. 2018), and a northeast Greenland fjord (Balmonte et al. 2020). Although their activities vary regionally, endo-acting enzymes produced by particle-associated microbial communities likely play a widespread role in efficient degradation of protein and carbohydrate constituents of particulate matter (Obayashi and Suzuki, 2005) from surface to deep waters (Fig. 7).

Latitudinal differences in particle-associated enzyme activities (Fig. 7) and the depths at which they sharply decline are robust, and may be linked to latitudinal differences in particle export and transfer efficiencies (Marsay et al. 2015; Weber et al. 2016; Henson et al. 2012; Henson et al. 2019). Low particle export efficiency at low latitudes (Henson et al. 2012) indicates intense remineralization in the epipelagic, consistent with the high and wide-ranging particle-associated enzymatic activities in our low latitude stations (Fig. 7). Such intense particulate matter degradation results in rapid particle flux attenuation (Marsay et al. 2015) and, thus, low particle transfer efficiencies (Weber et al. 2016), coincident with lower rates of particle-associated enzyme activities in low latitude deep waters (Fig. 5, S5b). In contrast, less intense remineralization in the epipelagic at high latitudes (Henson et al. 2012; Marsay et al. 2015) result in high particle export and transfer efficiencies to deep waters (Weber et al. 2016). High particle fluxes likely sustain the wide range of particle associated activities still detectable in the mesopelagic-bathypelagic transition at high latitudes (Fig. 5, S5b). That the highest total organic carbon concentration exported to sediments was measured in the northernmost station of the transect (Pohlner et al. 2017) indicates high particle export and transfer in the Bering Sea. As a caveat, recent findings demonstrate substantial differences in within-region particle export and transfer efficiencies (Henson et al. 2019). Nevertheless, we hypothesize that latitudinal differences in the enzymatic capabilities of particle-associated taxa (Fig. 7) may additionally influence variations in particle export and transfer efficiencies, consistent with observations of a primary role of particle fragmentation for particle fluxes (Briggs et al. 2020).

Finally, with increasing depth, peptidase and glucosidase activities became increasingly divergent between bulk water and particles (Fig. 3a, 7). This pattern bears a striking resemblance to that observed among bulk seawater and particle-associated bacterial community composition in the central Arctic (Balmonte et al. 2018). Microbial decision to remain particle-attached or detach can be predicted by the optimal foraging theory and patch use dynamics (Yawata et al. 2020). In particle-poor environments, such as the deep ocean, microbes increase residence time on particles to maximize fitness and avoid long search times for other particles. Reduced frequency of detachment would yield increasingly divergent particle-attached versus free-living taxa with increasing depth, with clear consequences in enzymatic patterns (Fig. 3a, 7). Longer particle residence time and distinct within-particle microbial community development trajectories (Thiele et al. 2015) could lead to large differences in particle-attached microbial community composition and activities in deep waters. Accordingly, variability in particle-associated enzyme activity patterns increased with increasing depth, but decreased for bulk water patterns (Figs. 3b, 7). These patterns are consistent with increased variability among deep water particle-associated microbial communities, due to regional differences in environmental conditions and particle quality and quantity (Salazar et al. 2016); such patterns are not evident for free-living taxa. A depth-related decrease in β-diversity in microbial metabolic potentials (Sunagawa et al. 2015) is consistent with enzymatic patterns in bulk water, but not on particles, reflecting the largely free-living lifestyle of microbes in the deep due to particle patchiness. Distinct bulk water and particle-associated enzyme patterns underscore the importance of particles in shaping microbial biogeochemical roles throughout the water column (Fig. 7).

## Conclusion

Latitudinal and depth differences in enzyme activities indicate substantial variations in microbial capabilities to degrade organic matter (Fig. 7). Enzyme-specific spatial trends and correlates, and different relative proportions of enzyme activities in bulk water and on particles, suggest varying sources, kinetics, and controls. Nevertheless, several features of enzyme activities can be generalized. Microbial communities employ narrower enzymatic spectra with increasing depth in bulk water and on particles, highlighting the importance of organic matter nature and quantity in structuring these patterns (Fig. 7). Moreover, activities of rarely-measured endopeptidases can exceed leucine aminopeptidase activities, especially on particles. Collectively, activities of a broad range of enzymes display latitudinal and depth-related trends in organic matter degradation consistent with emerging patterns of microbial community structure and function, and particles fluxes on a global scale.

## Acknowledgements

We thank the captain, crew, and scientific party of R/V Sonne for their excellent fieldwork support. Additionally, we thank Sherif Ghobrial, Karylle Abella-Hall, and Sarah Brown for assistance with sample processing. The cruise (SO248) was funded by the German Federal Ministry of Education and Research (BMBF) within the BacGeoPac project (03G0248A). JPB and CA were supported by NSF (OCE-1332881 and OCE-1736772 to CA). JPB was additionally funded by a UNC Dissertation Completion Fellowship, UNC Global Partnership Award, and a Carl Tryggers Postdoctoral Fellowship at Uppsala University. MS and HAG were also supported by Deutsche Forschungsgemeinschaft within the Collaborative Research Center Roseobacter (TRR51).

## References

Alderkamp, A.-C., M. Van Rijssel, and H. Bolhuis. 2007. Characterization of marine bacteria and the activity of their enzyme systems involved in degradation of the algal storage glucan laminarin. FEMS Microbiol. Ecol. 59: 108–117

Arnosti, C. 2003. Fluorescent derivatization of polysaccharides and carbohydrate-containing biopolymers for measurement of enzyme activities in complex media. J. Chromatogr. B. 793: 181–191

Arnosti, C. 2011. Microbial Extracellular Enzymes and the Marine Carbon Cycle. Annu. Rev. Mar. Sci. Vol 3. pp 401–425

Arnosti, C. 2020a. Microbial enzyme activities: glucosidase and peptidase activities of bulk seawater samples from the RV\Sonne cruise SO248 in the South and North Pacific, along 180 W, May, 2016. Biological and Chemical Oceanography Data Management Office (BCO-DMO). Dataset version 2018-07-31. doi:10.26008/1912/bco-dmo.743224.1 Accessed 2020-01-10

Arnosti, C. 2020b. Microbial enzyme activities: glucosidase and peptidase activities of gravity filtered seawater samples from the RV\Sonne cruise SO248 in the South and North Pacific, along 180 W, May, 2016. Biological and Chemical Oceanography Data Management Office (BCO-DMO). Dataset version 2018-07-31. doi:10.26008/1912/bco-dmo.743320.1 Accessed 2020-01-10

Arnosti, C. 2020c. Microbial enzyme activities: polysaccharide hydrolase activities in bulk seawater samples from the RV\Sonne cruise SO248 in the South and North Pacific, along 180 W, May, 2016. Biological and Chemical Oceanography Data Management Office (BCO-DMO). Dataset version 2018-07-31. doi:10.26008/1912/bco-dmo.743054.1 Accessed 2020-01-10

Arnosti, C. 2020d. Microbial enzyme activities: polysaccharide hydrolase activities of gravity filtered seawater samples from the R/V Sonne cruise SO248 in the South and North Pacific, along 180 W, May, 2016. Biological and Chemical Oceanography Data Management Office (BCO-DMO). Dataset version 2018-07-31. doi:10.26008/1912/bco-dmo.743274.1 Accessed 2020-01-10

Arnosti, C. 2015. Contrasting strategies in microbial degradation of organic matter in the water column and sediments: An example from Arctic fjords of Svalbard. Mar. Chem. 168: 151–156

Arnosti, C., A.D. Steen, K. Ziervogel, S. Ghobrial, and W.H. Jeffrey. 2011. Latitudinal Gradients in Degradation of Marine Dissolved Organic Carbon. PLoS ONE 6: e28900

Azam, F. and F. Malfatti. 2007. Microbial structuring of marine ecosystems (vol 5, pg 782-791, 2007). Nature Rev. Microbiol. 5: 966–U923.

Badewien, T.H., H. Winkler, K.L. Arndt, and M. Simon. 2016. Physical oceanography during SONNE cruise SO248 (BacGeoPac). Institute for Chemistry and Biology of the Marine Environment, Carl-von-Ossietzky University of Oldenburg, Germany, PANGAEA, https://doi.org/10.1594/PANGAEA.864673

Balmonte, J.P., A. Teske, and C. Arnosti. 2018. Structure and function of high Arctic pelagic, particle-associated and benthic bacterial communities. Environ. Microbiol. 20: 2941–2954.

Balmonte, J.P., A. Buckley, A. Hoarfrost, S. Ghobrial, K. Ziervogel, A. Teske, and others. 2019. Community structural differences shape microbial responses to high molecular weight organic matter. Environ. Microbiol. 21: 557–571.

Balmonte, J.P., H. Hasler-Sheetal, R.N. Glud, T.J. Andersen, M.K. Sejr, M. Middelboe, and others. 2020. Sharp contrasts between freshwater and marine microbial enzymatic capabilities, community composition, and DOM pools in a NE Greenland fjord. Limnol. Oceanogr. 65: 77–95.

Baltar, F., J. Aristegui, E. Sintes, H.M. van Aken, J.M. Gasol, and G.J. Herndl. 2009. Prokaryotic extracellular enzymatic activity in relation to biomass production and respiration in the meso- and bathypelagic waters of the (sub)tropical Atlantic. Environ. Microbiol. 11: 1998–2014.

Baltar, F., J. Arístegui, J.M. Gasol, E. Sintes, H.M. van Aken, and G.J. Herndl. 2010. High dissolved extracellular enzymatic activity in the deep central Atlantic Ocean. Aquat. Microb. Ecol. 58: 287–302.

Behrenfeld, M.J., R.T. O’Malley, D.A. Siegel, C.R. McClain, J.L. Sarmiento, G.C. Feldman, and others. 2006. Climate-driven trends in contemporary ocean productivity. Nature 444: 752–755.

Benner, R. 2002. Chemical Composition and Reactivity: p. 59–90. In D. A. Hansell and C. A. Carlson [eds.], Biogeochemistry of marine dissolved organic matter. Elsevier.

Bergauer, K., A. Fernandez-Guerra, J.A.L. Garcia, R.R. Sprenger, R. Stepanauskas, and M.G. Pachiadaki. 2018. Organic matter processing by microbial communities throughout the Atlantic water column as revealed by metaproteomics. Proc. Natl. Acad. Sci. USA 115: E400–E408.

Berlemont, R and A.C. Martiny. 2016. Glycoside hydrolases across environmental microbial communities. PLoS Comp. Biol. 12: e1005300.

Bong C.W., Y. Obayashi, and S. Suzuki. 2013. Succession of protease activity in seawater and bacterial isolates during starvation in a mesocosm experiment. Aquat. Microb. Ecol. 69: 33–46

Briggs, N, G. Dall’Olmo, and H. Claustre. 2020. Major role of particle fragmentation in regulating biological sequestration of CO2 by the oceans. Science 367: 791–793.

Christian, J.R. and D.M. Karl. 1995. Bacterial ectoenzymes in marine waters: Activity ratios and temperature responses in three oceanographic provinces. Limnol. Oceanogr. 40: 1042–1049.

D’Ambrosio, L., K. Ziervogel, B. MacGregor, A. Teske, and C. Arnosti. 2014. Composition and enzymatic function of particle-associated and free-living bacteria: a coastal/offshore comparison. ISME J. 8: 2167–2179.

Davey, K.E., R.R. Kirby, C.M. Turley, A.J. Weightman, and J.C. Fry. 2001. Depth variation of bacterial extracellular enzyme activity and population diversity in the northeastern North Atlantic Ocean. Deep Sea Res. Part II Top. Stud. Oceanogr. 48: 1003–1017.

Elifantz, H., L.A. Waidner, M.T. Cottrell, and D.L. Kirchman. 2008. Diversity and abundance of glycosyl hydrolase family 5 in the North Atlantic Ocean. FEMS Microbiol. Ecol. 63: 316–327.

Fuhrman, J.A., J.A. Steele, I. Hewson, M.S. Schwalbach, M.V. Brown, and J.L. Green. 2008. A latitudinal diversity gradient in planktonic marine bacteria. Proc. Natl. Acad. Sci. USA. 105: 7774–7778.

Fukuda, R, Y. Sohrin, N. Saotome, H. Fukuda, T. Nagata, and I. Koike. 2000. East—west gradient in ectoenzyme activities in the subarctic Pacific: Possible regulation by zinc. Limnol. Oceanogr. 45: 930–939.

Ghiglione, J.F., P.E. Galand, T. Pommier, C. Pedros-Alio, E.W. Maas, K. Bakker, and others. 2012. Pole-to-pole biogeography of surface and deep marine bacterial communities. Proc. Natl. Acad. Sci. USA. 109: 17633–17638.

Giebel, H.A., M. Wolterink, T. Brinkhoff, and M. Simon (2019). Complementary energy acquisition via aerobic anoxygenic photosynthesis and carbon monoxide oxidation by Planktomarina temperata of the Roseobacter group. FEMS Microbiology Ecology 95(5): fiz050.

Giebel, H.A, K.L. Arndt KL, C. Arnosti, T.H. Badewien, I. Bakenhus, J.P. Balmonte, and others. 2020: Hydrography, Biogeochemistry, microbial population, growth and substrate dynamics between subarctic and subantarctic waters in the Pacific Ocean during the cruises SO248 and SO254 with RV Sonne. PANGAEA, https://doi.pangaea.de/10.1594/PANGAEA.918500

Henson, S., F. Le Moigne, and S. Giering. 2019. Drivers of Carbon Export Efficiency in the Global Ocean. Global Biogeochem. Cycles 33: 891–903.

Henson, S.A., R. Sanders, and E. Madsen. 2012. Global patterns in efficiency of particulate organic carbon export and transfer to the deep ocean. Global Biogeochem. Cycles 26.

Hoarfrost, A. and C. Arnosti. 2017. Heterotrophic Extracellular Enzymatic Activities in the Atlantic Ocean Follow Patterns Across Spatial and Depth Regimes. Front. Mar. Sci. 4.

Ibarbalz, F.M., N. Henry, M.C. Brandão, S. Martini, G. Busseni, H. Byrne, and others. 2019. Global Trends in Marine Plankton Diversity across Kingdoms of Life. Cell 179: 1084–1097.e1021.

Ladau J, T.J. Sharpton, M.M. Finucane, G. Jospin, S.W. Kembel, J. O’Dwyer, and others. 2013. Global marine bacterial diversity peaks at high latitudes in winter. ISME J. 7: 1669–1677.

Louca, S., L.W. Parfrey, and M. Doebeli. 2016. Decoupling function and taxonomy in the global ocean microbiome. Science 353: 1272–1277.

Lunau, M., A. Lemke, O. Dellwig, and M. Simon (2006). Physical and biogeochemical controls of microaggregate dynamics in a tidally affected coastal ecosystem. Limnol. Oceanogr. 51: 847–859.

Marsay, C.M., R.J. Sanders, S.A. Henson, K. Pabortsava, E.P. Achterberg, R.S. Lampitt. 2015. Attenuation of sinking particulate organic carbon flux through the mesopelagic ocean. Proc. Natl. Acad. Sci. USA. 112: 1089–1094.

Neumann, A.M., J.P. Balmonte, M. Berger, H.A. Giebel, C. Arnosti, S. Voget and others. 2015. Different utilization of alginate and other algal polysaccharides by marine Alteromonas macleodii ecotypes. Environ. Microbiol. 10: 3857–3868

Obayashi, Y. and S. Suzuki. 2005. Proteolytic enzymes in coastal surface seawater: Significant activity of endopeptidases and exopeptidases. Limnol. Oceanogr. 50: 722–726.

Painter, T.J. 1983. Algal Polysaccharides: p. 195–285. In G. O. Aspinall [ed], The polysaccharides, v. 2. Academic Press.

Pohlner, M., J. Degenhardt, A.J.E. von Hoyningen-Huene, B. Wemheuer, N. Erlmann, B. Schnetger, and others. 2017. The Biogeographical Distribution of Benthic Roseobacter Group Members along a Pacific Transect Is Structured by Nutrient Availability within the Sediments and Primary Production in Different Oceanic Provinces. Front. Microbiol. 8.

Pommier, T., B. Canbäck, L. Riemann, K.H. Bostrom, K. Simu, P. Lundberg, and others. 2007. Global patterns of diversity and community structure in marine bacterioplankton. Molec. Ecol. 16: 867–880.

Raes, E.J., L. Bodrossy, J. van de Kamp, A. Bissett, M. Ostrowski, M.V. Brown, and others. 2018. Oceanographic boundaries constrain microbial diversity gradients in the South Pacific Ocean. Proc. Natl. Acad. Sci. USA. 115: E8266–E8275.

Raes, J., I. Letunic, T. Yamada, L.J. Jensen, and P. Bork. 2011. Toward molecular trait-based ecology through integration of biogeochemical, geographical and metagenomic data. Molec. Syst. Biol. 7.

Salazar, G., F.M. Cornejo-Castillo, V. Benítez-Barrios, E. Fraile-Nuez, X.A. Álvarez-Salgado, C.M. Duarte, and others. 2016. Global diversity and biogeography of deep-sea pelagic prokaryotes. ISME J. 10: 596–608.

Salazar, G., L. Paoli, A. Alberti, J. Huerta-Cepas, H.J. Ruscheweyh, M. Cuenca, and others. 2019. Gene Expression Changes and Community Turnover Differentially Shape the Global Ocean Metatranscriptome. Cell 179: 1068–1083.e1021.

Simon, M. and F. Azam. 1989. Protein content and protein synthesis rates of planktonic marine bacteria. Mar. Ecol. Prog. Ser. 51: 201–213.

Simon, M, B. Rosenstock, and W. Zwisler. 2004. Coupling of epipelagic and mesopelagic heterotrophic picoplankton production to phytoplankton biomass in the Antarctic polar frontal region. Limnol. Oceanogr. 49: 1035–1043.

Smith, D.C., M. Simon, A.L. Alldredge, and F. Azam. 1992. Intense hydrolytic enzyme activity on marine aggregates and implications for rapid particle dissolution. Nature 359: 139–142.

Steen, A.D., K. Ziervogel, S. Ghobrial, and C. Arnosti. 2012. Functional variation among polysaccharide-hydrolyzing microbial communities in the Gulf of Mexico. Mar. Chem. 138: 13–20.

Steen A.D., and C. Arnosti. 2013. Extracellular peptidase and carbohydrate hydrolase activities in an Arctic fjord (Smeerenburgfjord, Svalbard). Aquat. Microb. Ecol. 69: 93–99.

Sul, W.J., T.A. Oliver, H.W. Ducklow, L.A. Amaral-Zettler, and M.L. Sogin. 2013. Marine bacteria exhibit a bipolar distribution. Proc. Natl. Acad. Sci. USA. 110: 2342–2347.

Sunagawa, S., L.P. Coelho, S. Chaffron, J.R. Kultima, K. Labadie, G. Salazar G, and others. 2015. Structure and function of the global ocean microbiome. Science 348.

Teeling, H., B.M. Fuchs, D. Becher, C. Klockow, A. Gardebrecht, C.M. Bennke, and others. 2012. Substrate-controlled succession of marine bacterioplankton populations induced by a phytoplankton bloom. Science 336: 608–611.

Teeling, H., B.M. Fuchs, C.M. Bennke, K. Krüger, M. Chafee, L. Kappelmann, and others. 2016. Recurring patterns in bacterioplankton dynamics during coastal spring algae blooms. eLife 5: e11888.

Teske, A., A. Durbin, K. Ziervogel, C. Cox, and C. Arnosti. 2011. Microbial Community Composition and Function in Permanently Cold Seawater and Sediments from an Arctic Fjord of Svalbard. Appl. Environ. Microbiol. 77: 2008–2018.

Thiele, S., B.M. Fuchs, R. Amann, and M.H. Iversen. 2015. Colonization in the Photic Zone and Subsequent Changes during Sinking Determine Bacterial Community Composition in Marine Snow. Appl. Environ. Microbiol. 81: 1463–1471.

Vetter, Y.A., J.W. Deming, P.A. Jumars, and B.B. Krieger-Brockett. 1998. A Predictive Model of Bacterial Foraging by Means of Freely Released Extracellular Enzymes. Microb. Ecol. 36: 75–92.

Weber, T., J.A. Cram, S.W. Leung, T. DeVries, and C. Deutsch. 2016. Deep ocean nutrients imply large latitudinal variation in particle transfer efficiency. Proc. Natl. Acad. Sci. USA. 113: 8606–8611.

Weiss, M.S., U. Abele, J. Weckesser, W. Welte, E. Schiltz, and G.E. Schulz. 1991. Molecular architecture and electrostatic properties of a bacterial porin. Science 254: 1627–1630.

Wietz, M., L. Gram, B. Jørgensen, and A. Schramm (2010). Latitudinal patterns in the abundance of major marine bacterioplankton groups. Aquat. Microb. Ecol. 61: 179.

Zaccone, R., L.S Monticelli, A. Seritti, C. Santinelli, M. Azzaro, A. Boldrin, and others. 2003. Bacterial processes in the intermediate and deep layers of the Ionian Sea in winter 1999: Vertical profiles and their relationship to the different water masses. J. Geophys. Res. Oceans 108.

Zhao, Z, F. Baltar, and G.J. Herndl. 2020. Linking extracellular enzymes to phylogeny indicates a predominantly particle-associated lifestyle of deep-sea prokaryotes. Sci. Adv. 6:eaaz4354

Ziervogel, K. and C. Arnosti. 2020. Substantial Carbohydrate Hydrolase Activities in the Water Column of the Guaymas Basin (Gulf of California). Front. Mar. Sci. 6.

