## Supplementary Information for "A Sea Change in Microbial Enzymes: Heterogeneous latitudinal and depth-related gradients in bulk water and particle-associated enzymatic activities from 30°S to 59°N in the Pacific Ocean"

| Stn. | Depth (m) | Depth layer | Date | CTD Event | Lat. | Long. | Temp. (°C) | Sal. (PSU) | Fluor. (µg/L) |
|---|---|---|---|---|---|---|---|---|---|
| S01 | 20 | Epipelagic (surface) | 2-May-16 | SO248_01-1 | 30.00°S | 177.00°E | 23.5 | 35.92 | 0.07 |
| S01 | 100 | Epipelagic (DCM) | 2-May-16 | SO248_01-1 | 30.00°S | 177.00°E | 19.5 | 35.86 | 0.50 |
| S01 | 300 | Mesopelagic (300m) | 2-May-16 | SO248_01-1 | 30.00°S | 177.00°E | 15.1 | 35.50 | 0.01 |
| S01 | 500 | Mesopelagic (500m) | 2-May-16 | SO248_01-1 | 30.00°S | 177.00°E | 11.3 | 35.01 | 0.01 |
| S01 | 1000 | Bathypelagic (1000m) | 2-May-16 | SO248_01-1 | 30.00°S | 177.00°E | 5.4 | 34.49 | 0.01 |
| S02 | 20 | Epipelagic (surface) | 3-May-16 | SO248_02-1 | 26.99°S | 178.21°E | 25.2 | 35.64 | 0.15 |
| S02 | 100 | Epipelagic (DCM) | 3-May-16 | SO248_02-1 | 26.99°S | 178.21°E | 21.5 | 35.88 | 0.44 |
| S02 | 300 | Mesopelagic (300m) | 3-May-16 | SO248_02-1 | 26.99°S | 178.21°E | 16.1 | 35.54 | 0.01 |
| S02 | 500 | Mesopelagic (500m) | 3-May-16 | SO248_02-1 | 26.99°S | 178.21°E | 11.5 | 35.05 | 0.01 |
| S02 | 1000 | Bathypelagic (1000m) | 3-May-16 | SO248_02-1 | 26.99°S | 178.21°E | 5.0 | 34.50 | 0.01 |
| S03 | 20 | Epipelagic (surface) | 6-May-16 | SO248_03-1 | 15.00°S | 178.00°W | 28.7 | 35.17 | 0.03 |
| S03 | 100 | Epipelagic (DCM) | 6-May-16 | SO248_03-1 | 15.00°S | 178.00°W | 24.0 | 36.19 | 0.51 |
| S04 | 20 | Epipelagic (surface) | 8-May-16 | SO248_04-3 | 10.33°S | 176.48°W | 29.9 | 34.69 | 0.07 |
| S04 | 90 | Epipelagic (DCM) | 8-May-16 | SO248_04-3 | 10.33°S | 176.48°W | 27.4 | 36.04 | 0.90 |
| S04 | 300 | Mesopelagic (300m) | 8-May-16 | SO248_04-3 | 10.33°S | 176.48°W | 13.2 | 35.07 | 0.02 |
| S04 | 1000 | Bathypelagic (1000m) | 8-May-16 | SO248_04-3 | 10.33°S | 176.48°W | 4.4 | 34.66 | 0.02 |
| S04 | 4000 | Bathypelagic (bottom) | 8-May-16 | SO248_04-3 | 10.33°S | 176.48°W | 1.0 | 34.84 | 0.02 |
| S05 | 20 | Epipelagic (surface) | 9-May-16 | SO248_05-1 | 5.00°S | 178.32°W | 30.4 | 34.56 | 0.12 |
| S05 | 90 | Epipelagic (DCM) | 9-May-16 | SO248_05-1 | 5.00°S | 178.32°W | 29.0 | 35.54 | 0.91 |
| S06 | 20 | Epipelagic (surface) | 11-May-16 | SO248_06-2 | 0.00 | 180.00 | 29.4 | 35.19 | 0.81 |
| S06 | 60 | Epipelagic (DCM) | 11-May-16 | SO248_06-2 | 0.00 | 180.00 | 28.9 | 35.24 | 1.15 |
| S06 | 300 | Mesopelagic (300m) | 11-May-16 | SO248_06-2 | 0.00 | 180.00 | 11.2 | 34.94 | 0.03 |
| S06 | 1000 | Bathypelagic (1000m) | 11-May-16 | SO248_06-2 | 0.00 | 180.00 | 4.3 | 34.69 | 0.04 |
| S06 | 4000 | Bathypelagic (bottom) | 11-May-16 | SO248_06-2 | 0.00 | 180.00 | 1.1 | 34.83 | 0.02 |
| S07 | 20 | Epipelagic (surface) | 13-May-16 | SO248_07-1 | 4.66°N | 179.40°E | 28.8 | 34.32 | 0.05 |
| S07 | 75 | Epipelagic (75m) | 13-May-16 | SO248_07-1 | 4.66°N | 179.40°E | 28.6 | 34.45 | 0.28 |
| S07 | 105 | Epipelagic (DCM) | 13-May-16 | SO248_07-1 | 4.66°N | 179.40°E | 24.4 | 34.93 | 0.63 |
| S07 | 300 | Mesopelagic (300m) | 13-May-16 | SO248_07-1 | 4.66°N | 179.40°E | 9.3 | 34.77 | 0.05 |
| S07 | 1000 | Bathypelagic (1000m) | 13-May-16 | SO248_07-1 | 4.6661°N | 179.40°E | 4.4 | 34.70 | 0.05 |
| S08 | 20 | Epipelagic (surface) | 15-May-16 | SO248_08-4 | 11.00°N | 179.00°E | 27.8 | 34.60 | 0.03 |
| S08 | 120 | Epipelagic (DCM) | 15-May-16 | SO248_08-4 | 11.00°N | 179.00°E | 21.7 | 34.82 | 0.87 |
| S09 | 20 | Epipelagic (surface) | 17-May-16 | SO248_09-6 | 16.00°N | 179.00°E | 26.6 | 35.09 | 0.02 |
| S09 | 120 | Epipelagic (DCM) | 17-May-16 | SO248_09-6 | 16.00°N | 179.00°E | 22.8 | 35.44 | 0.48 |
| S10 | 20 | Epipelagic (surface) | 18-May-16 | SO248_10-2a | 22.00°N | 178.32°E | 25.0 | 35.52 | 0.07 |
| S10 | 120 | Epipelagic (DCM) | 18-May-16 | SO248_10-2a | 22.00°N | 178.32°E | 19.5 | 35.21 | 0.63 |
| S10 | 300 | Mesopelagic (300m) | 18-May-16 | SO248_10-2a | 22.00°N | 178.32°E | 13.1 | 34.49 | 0.01 |
| S10 | 1000 | Bathypelagic (1000m) | 18-May-16 | SO248_10-2a | 22.00°N | 178.32°E | 3.8 | 34.60 | 0.03 |
| S10 | 2803 | Bathypelagic (bottom) | 18-May-16 | SO248_10-2a | 22.00°N | 178.32°E | 1.4 | 34.81 | 0.02 |
| S11 | 20 | Epipelagic (surface) | 20-May-16 | SO248_11-1 | 28.00°N | 177.33°E | 20.8 | 35.08 | 0.06 |
| S11 | 70 | Epipelagic (DCM) | 20-May-16 | SO248_11-1 | 28.00°N | 177.33°E | 17.5 | 34.90 | 1.52 |
| S12 | 20 | Epipelagic (surface) | 21-May-16 | SO248_12-1 | 34.00°N | 177.33°E | 15.7 | 34.76 | 1.05 |
| S12 | 50 | Epipelagic (DCM) | 21-May-16 | SO248_12-1 | 34.00°N | 177.33°E | 15.5 | 34.75 | 1.40 |
| S13 | 20 | Epipelagic (surface) | 23-May-16 | SO248_13-3 | 39.97°N | 177.33°E | 11.6 | 34.33 | 3.99 |
| S13 | 50 | Epipelagic (DCM) | 23-May-16 | SO248_13-3 | 39.97°N | 177.33°E | 11.2 | 34.32 | 2.24 |
| S14 | 20 | Epipelagic (surface) | 24-May-16 | SO248_14-3 | 45.00°N | 178.75°E | 5.9 | 33.31 | 1.87 |
| S14 | 60 | Epipelagic (DCM) | 24-May-16 | SO248_14-3 | 45.00°N | 178.75°E | 5.7 | 33.38 | 0.56 |
| S14 | 300 | Mesopelagic (300m) | 24-May-16 | SO248_14-3 | 45.00°N | 178.75°E | 4.5 | 33.96 | 0.04 |
| S14 | 1000 | Bathypelagic (1000m) | 24-May-16 | SO248_14-3 | 45.00°N | 178.75°E | 2.9 | 34.52 | 0.06 |
| S14 | 4000 | Bathypelagic (bottom) | 24-May-16 | SO248_14-3 | 45.00°N | 178.75°E | 1.2 | 34.83 | 0.03 |
| S15 | 20 | Epipelagic (surface) | 25-May-16 | SO248_15-1 | 47.50°N | 179.13°E | 4.5 | 33.09 | 1.68 |

| | | | | | | | | | |
|---|---|---|---|---|---|---|---|---|---|
| S15 | 60 | Epipelagic (DCM) | 25-May-16 | SO248_15-1 | 47.50°N | 179.13°E | 3.5 | 33.11 | 0.79 |
| S16 | 20 | Epipelagic (surface) | 26-May-16 | SO248_16-2 | 50.00°N | 179.55°E | 4.8 | 33.07 | 1.55 |
| S16 | 60 | Epipelagic (DCM) | 26-May-16 | SO248_16-2 | 50.00°N | 179.55°E | 3.4 | 33.13 | 0.80 |
| S16 | 300 | Mesopelagic (300m) | 26-May-16 | SO248_16-2 | 50.00°N | 179.55°E | 3.8 | 34.20 | 0.07 |
| S16 | 1000 | Bathypelagic (1000m) | 26-May-16 | SO248_16-2 | 50.00°N | 179.55°E | 2.6 | 34.57 | 0.05 |
| S16 | 4000 | Bathypelagic (bottom) | 26-May-16 | SO248_16-2 | 50.00°N | 179.55°E | 1.1 | 34.84 | 0.02 |
| S17 | 20 | Epipelagic (surface) | 28-May-16 | SO248_17-4 | 54.00°N | 179.58°E | 4.6 | 33.20 | 1.47 |
| S17 | 60 | Epipelagic (DCM) | 28-May-16 | SO248_17-4 | 54.00°N | 179.58°E | 2.8 | 33.24 | 0.56 |
| S18 | 20 | Epipelagic (surface) | 29-May-16 | SO248_18-3 | 57.00°N | 179.58°E | 4.2 | 33.17 | 1.03 |
| S18 | 60 | Epipelagic (DCM) | 29-May-16 | SO248_18-3 | 57.00°N | 179.58°E | 2.5 | 33.22 | 0.36 |
| S18 | 300 | Mesopelagic (300m) | 29-May-16 | SO248_18-3 | 57.00°N | 179.58°E | 3.9 | 33.98 | 0.06 |
| S18 | 1000 | Bathypelagic (1000m) | 29-May-16 | SO248_18-3 | 57.00°N | 179.58°E | 2.8 | 34.51 | 0.05 |
| S18 | 3500 | Bathypelagic (bottom) | 29-May-16 | SO248_18-3 | 57.00°N | 179.58°E | 1.3 | 34.82 | 0.04 |
| S19 | 20 | Epipelagic (surface) | 30-May-16 | SO248_19-1 | 58.90°N | 179.00°W | 5.4 | 32.90 | 1.86 |
| S19 | 60 | Epipelagic (DCM) | 30-May-16 | SO248_19-1 | 58.90°N | 179.00°W | 4.1 | 33.05 | 0.62 |

**Table S1.** List of all stations and depths. Rows with gray shading denote stations for which only bulk peptidase and glucosidase activities were measured at the surface and DCM; there are no data from below the DCM. Rows with no shading denote the main stations. Stn = Station; Lat = Latitude; Long = Longitude; Temp = Temperature; Sal = Salinity; Fluor = Fluorescence; surf = surface; DCM = deep chlorophyll maximum. Temperature, salinity, and fluorescence values were obtained from the shipboard CTD.

| Stn. | Depth (m) | Depth layer | Date | Filter 1 volume (L) | Filter 2 volume (L) | Filter 3 volume (L) |
|---|---|---|---|---|---|---|
| S01 | 100 | Epipelagic (DCM) | 2-May-16 | 6.88 | 7.76 | 7.85 |
| S01 | 300 | Mesopelagic | 2-May-16 | 7.70 | 8.15 | 7.32 |
| S01 | 1000 | Bathypelagic | 2-May-16 | 6.00 | 7.85 | 7.72 |
| S02 | 100 | Epipelagic (DCM) | 3-May-16 | 7.22 | 7.66 | 7.68 |
| S02 | 300 | Mesopelagic | 3-May-16 | 7.78 | 7.02 | 7.57 |
| S02 | 1000 | Bathypelagic | 3-May-16 | 8.00 | 7.83 | 7.78 |
| S04 | 90 | Epipelagic (DCM) | 8-May-16 | 6.05 | 7.72 | 7.05 |
| S04 | 300 | Mesopelagic | 8-May-16 | 7.90 | 7.88 | 7.74 |
| S04 | 1000 | Bathypelagic | 8-May-16 | 8.05 | 7.55 | 7.73 |
| S06 | 60 | Epipelagic (DCM) | 11-May-16 | 7.42 | 7.12 | 6.34 |
| S06 | 300 | Mesopelagic | 11-May-16 | 7.30 | 7.55 | 7.45 |
| S06 | 1000 | Bathypelagic | 11-May-16 | 7.35 | 6.89 | 7.54 |
| S07 | 105 | Epipelagic (DCM) | 13-May-16 | 7.46 | 7.78 | 7.80 |
| S07 | 300 | Mesopelagic | 13-May-16 | 8.14 | 8.05 | 8.05 |
| S07 | 1000 | Bathypelagic | 13-May-16 | 7.66 | 7.75 | 7.89 |
| S10 | 120 | Epipelagic (DCM) | 18-May-16 | 7.80 | 6.90 | 6.30 |
| S10 | 300 | Mesopelagic | 18-May-16 | 7.23 | 7.82 | 7.74 |
| S10 | 1000 | Bathypelagic | 18-May-16 | 7.75 | 7.65 | 7.89 |
| S14 | 60 | Epipelagic (DCM) | 24-May-16 | 6.18 | 6.00 | 6.45 |
| S14 | 300 | Mesopelagic | 24-May-16 | 7.48 | 8.12 | 8.06 |
| S14 | 1000 | Bathypelagic | 24-May-16 | 7.75 | 7.80 | 7.84 |
| S16 | 60 | Epipelagic (DCM) | 26-May-16 | 6.14 | 7.57 | 6.90 |
| S16 | 300 | Mesopelagic | 26-May-16 | 8.15 | 8.10 | 8.10 |
| S16 | 1000 | Bathypelagic | 26-May-16 | 8.02 | 7.90 | 7.90 |
| S18 | 60 | Epipelagic (DCM) | 29-May-16 | 7.15 | 7.30 | 6.95 |
| S18 | 300 | Mesopelagic | 29-May-16 | 7.95 | 8.05 | 7.93 |
| S18 | 1000 | Bathypelagic | 29-May-16 | 7.78 | 7.90 | 8.00 |

**Table S2**. Volumes of water filtered for particle-associated enzyme activities. Stn = Station; Lat = Latitude; Long = Longitude; Temp = Temperature; Sal = Salinity; Fluor = Fluorescence; surf = surface; DCM = deep chlorophyll maximum.

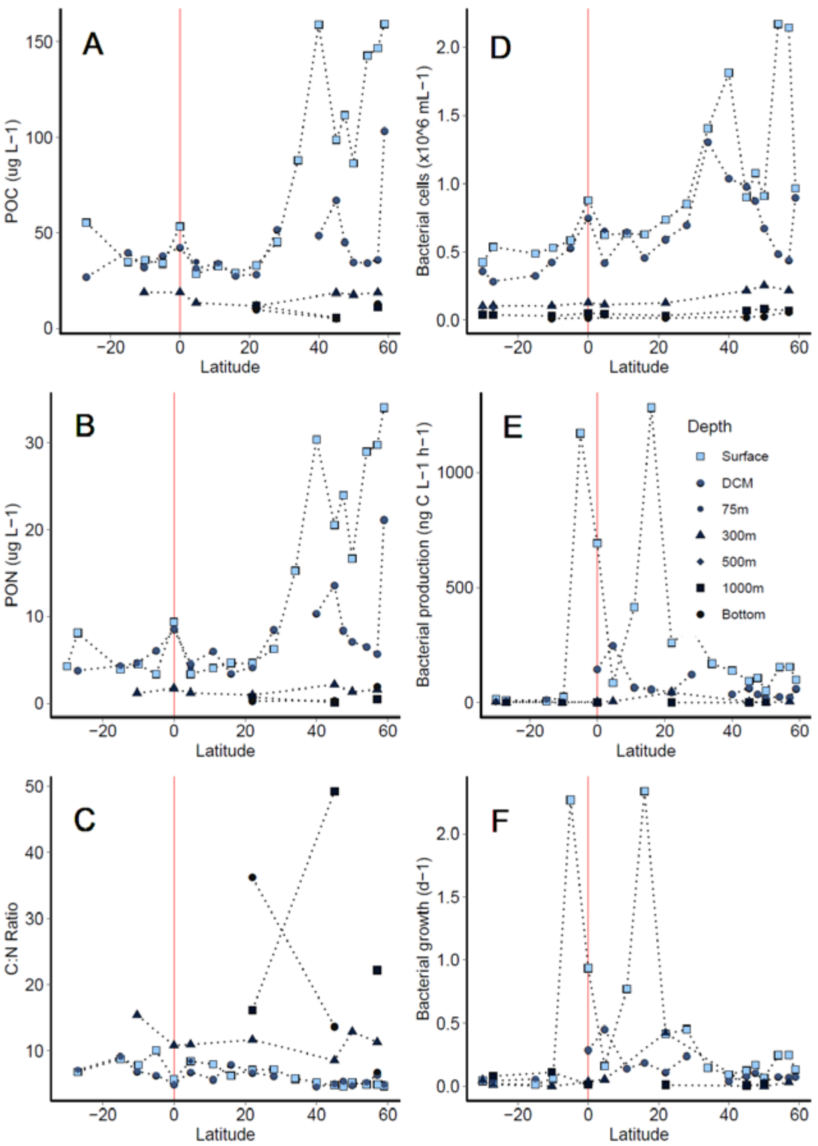


**Figure S1.** Biotic and abiotic parameters along the latitudinal transect. Negative latitudes indicate °S. Red vertical line denotes the equator. POC = particulate organic carbon concentrations (**a**), PON = particulate organic nitrogen concentrations (**b**), POC:PON ratios (**c**), Bacterial cell counts (**d**), bacterial production, (**e**), and bacterial growth rates (**f**). Note differences in units and y-axis scales.

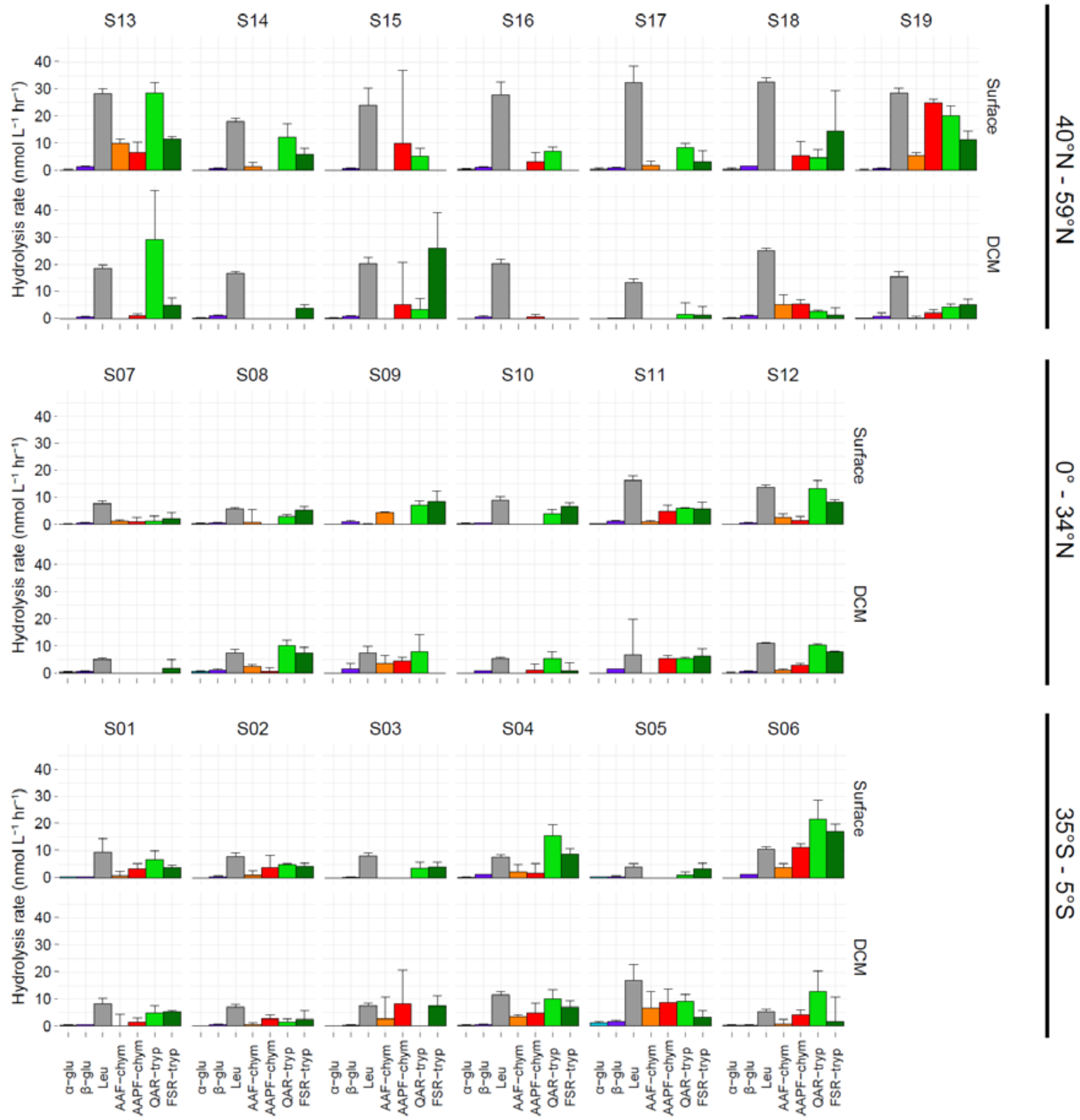


**Figure S2a.** Surface and deep chlorophyll maximum (DCM) bulk water (non-size fractionated) glucosidase and peptidase activities at all stations. Note that DCM depths vary by station (*see Table S1*), and x-axis scales differ by substrate. Error bars represent the standard deviation of rates from triplicate incubations. Rates shown were measured at 12 h. glu = glucosidase, leu = leucine aminopeptidase, chym = chymotrypsin, tryp = trypsin, AAF = alanine-alanine-phenylalanine, AAPF = alanine-alanine-proline-phenylalanine, QAR = glutamine-alanine-arginine, FSR = phenylalanine-serine-arginine.

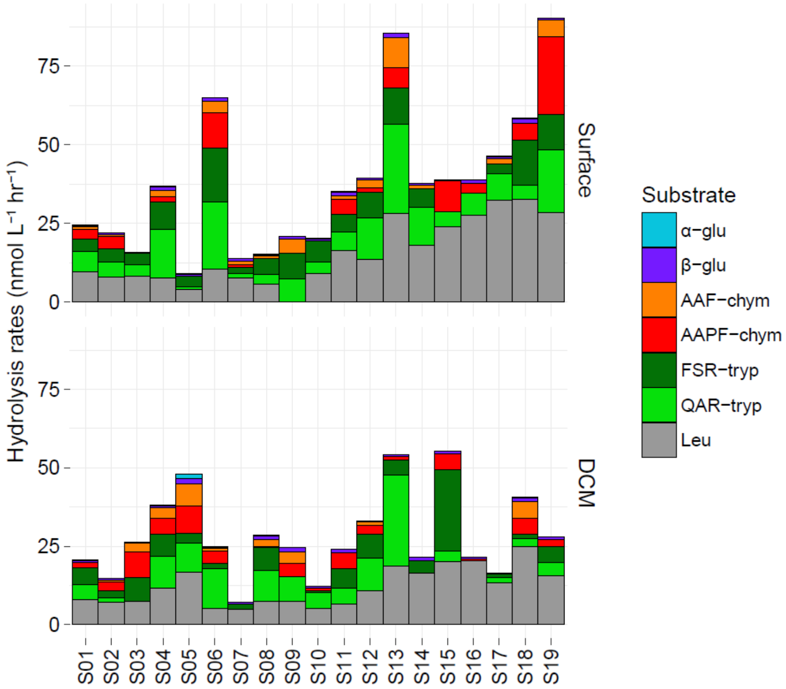


**Figure S2b.** Summed rates of surface and deep chlorophyll maximum (DCM) bulk water (non-size fractionated) glucosidase and peptidase activities at all stations. Note that DCM depths vary by station (*see Table S1*). Rates shown were measured at 12 h. glu = glucosidase, leu = leucine aminopeptidase, chym = chymotrypsin, tryp = trypsin, AAF = alanine-alanine-phenylalanine, AAPF = alanine-alanine-proline-phenylalanine, QAR = glutamine-alanine-arginine, FSR = phenylalanine-serine-arginine.

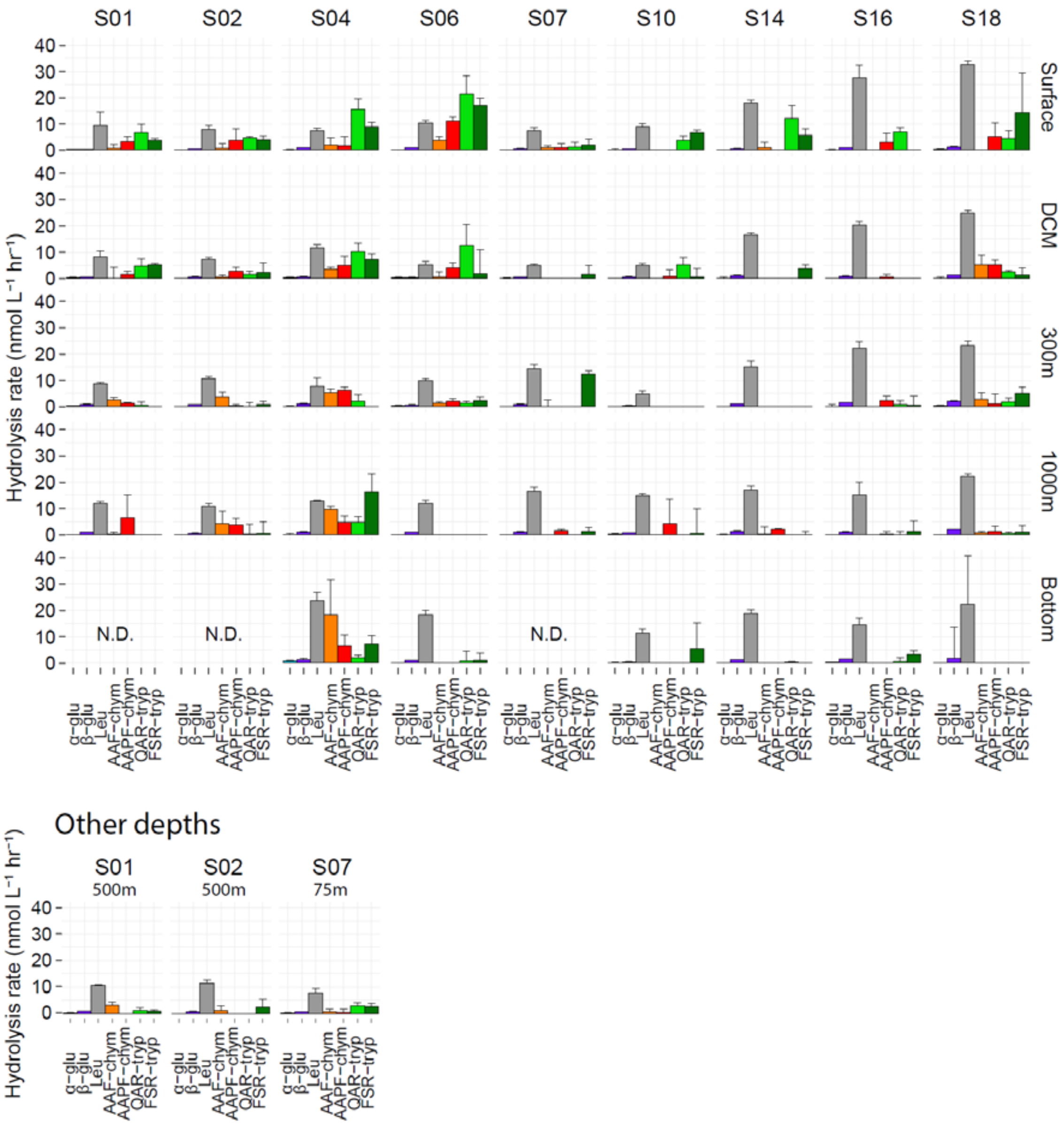


**Figure S3a.** Bulk water (non-size fractionated) glucosidase and peptidase activities at all depths at the main stations. Note that DCM and bottom water depths vary by station (*see Table S1*). Error bars represent the standard deviation of rates from triplicate incubations. Rates shown were measured at 12 h. glu = glucosidase, leu = leucine aminopeptidase, chym = chymotrypsin, tryp = trypsin, AAF = alanine-alanine-phenylalanine, AAPF = alanine-alanine-proline-phenylalanine, QAR = glutamine-alanine-arginine, FSR = phenylalanine-serine-arginine, N.D. = No Data collected.

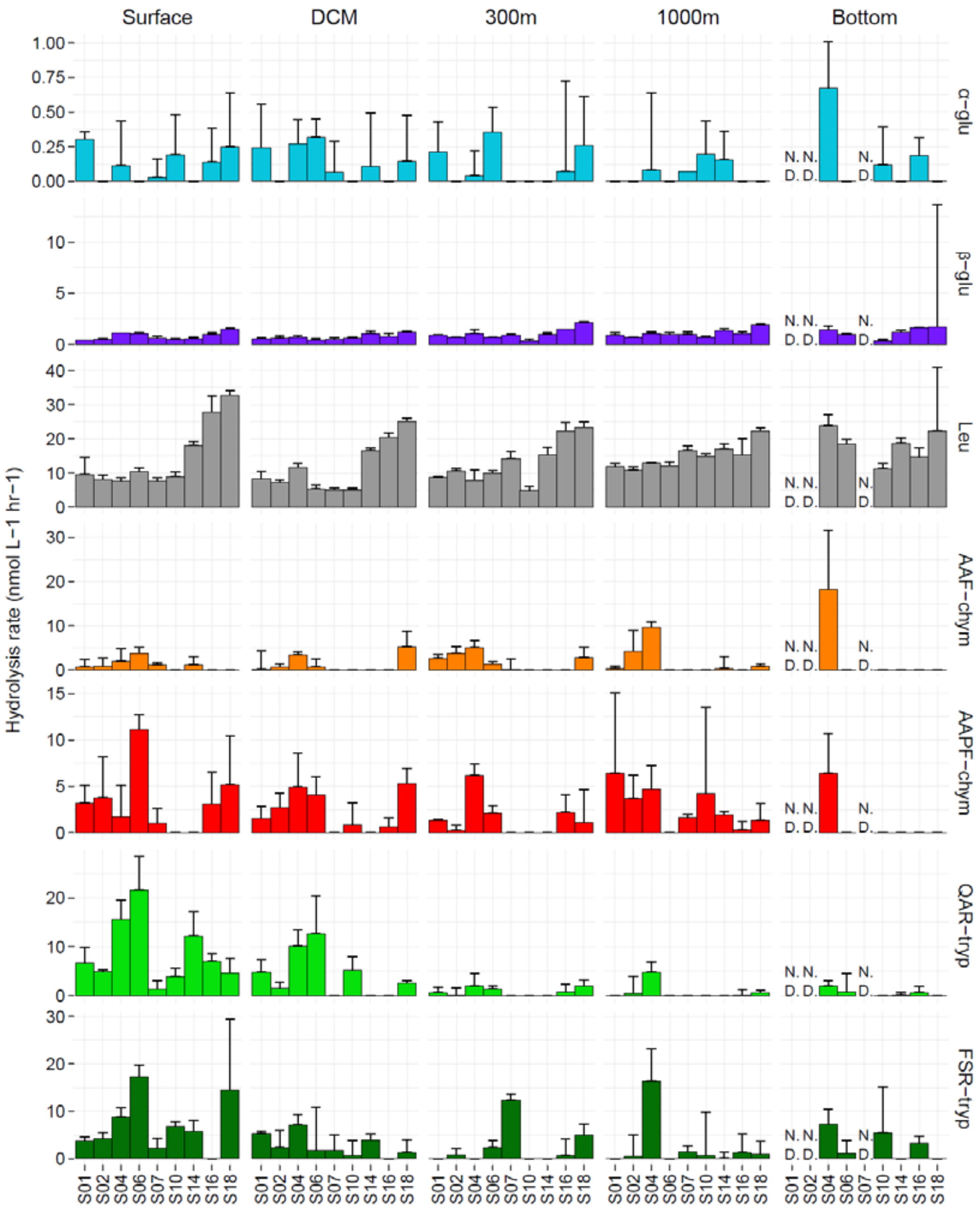


**Figure S3b.** Substrate-separated rates for bulk water (non-size fractionated) glucosidase and peptidase activities at all depths at the main stations. Note that DCM and bottom water depths vary by station (*see Table S1*). Error bars represent the standard deviation of rates from triplicate incubations. Rates shown were measured at 12 h. glu = glucosidase, leu = leucine aminopeptidase, chym = chymotrypsin, tryp = trypsin, AAF = alanine-alanine-phenylalanine, AAPF = alanine-alanine-proline-phenylalanine, QAR = glutamine-alanine-arginine, FSR = phenylalanine-serine-arginine, N.D. = No Data collected.

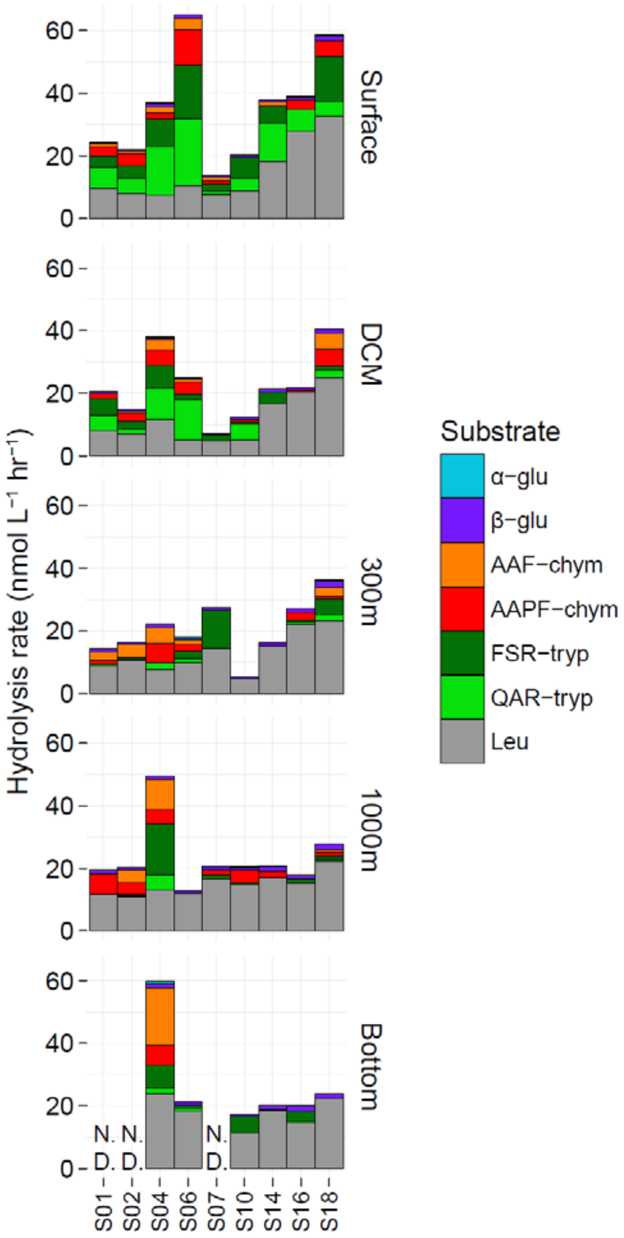


**Figure S3c.** Summed rates of bulk water (non-size fractionated) glucosidase and peptidase activities at all depths of the main stations. Note that DCM and bottom water depths vary by station (*see Table S1*). Rates shown were measured at 12 h. glu = glucosidase, leu = leucine aminopeptidase, chym = chymotrypsin, tryp = trypsin, AAF = alanine-alanine-phenylalanine, AAPF = alanine-alanine-proline-phenylalanine, QAR = glutamine-alanine-arginine, FSR = phenylalanine-serine-arginine, N.D. = No Data collected.

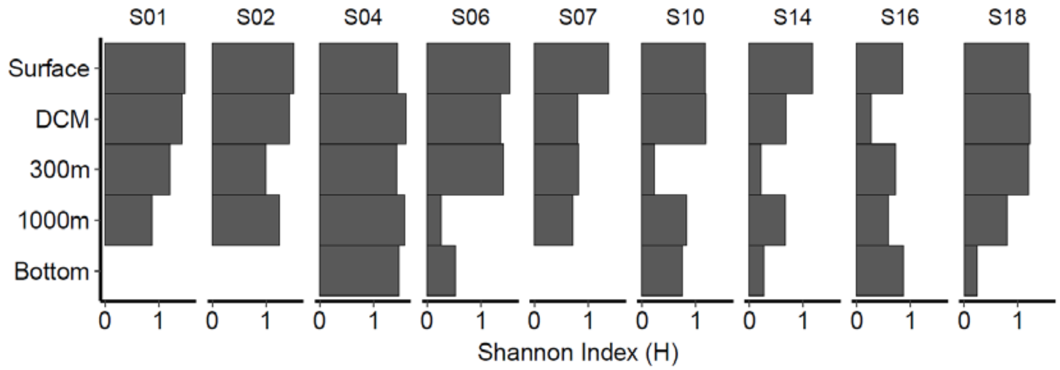


.

**Figure S3d.** Shannon indices for bulk water (non-size fractionated) glucosidase and peptidase activities at all depths at the main stations. Note that DCM and bottom water depths vary by station (*see Table S1*). Shannon indices were calculated from rates measured at 12 h, based on a previously established equation (Steen *et al.*, 2010). N.D. = No Data collected.

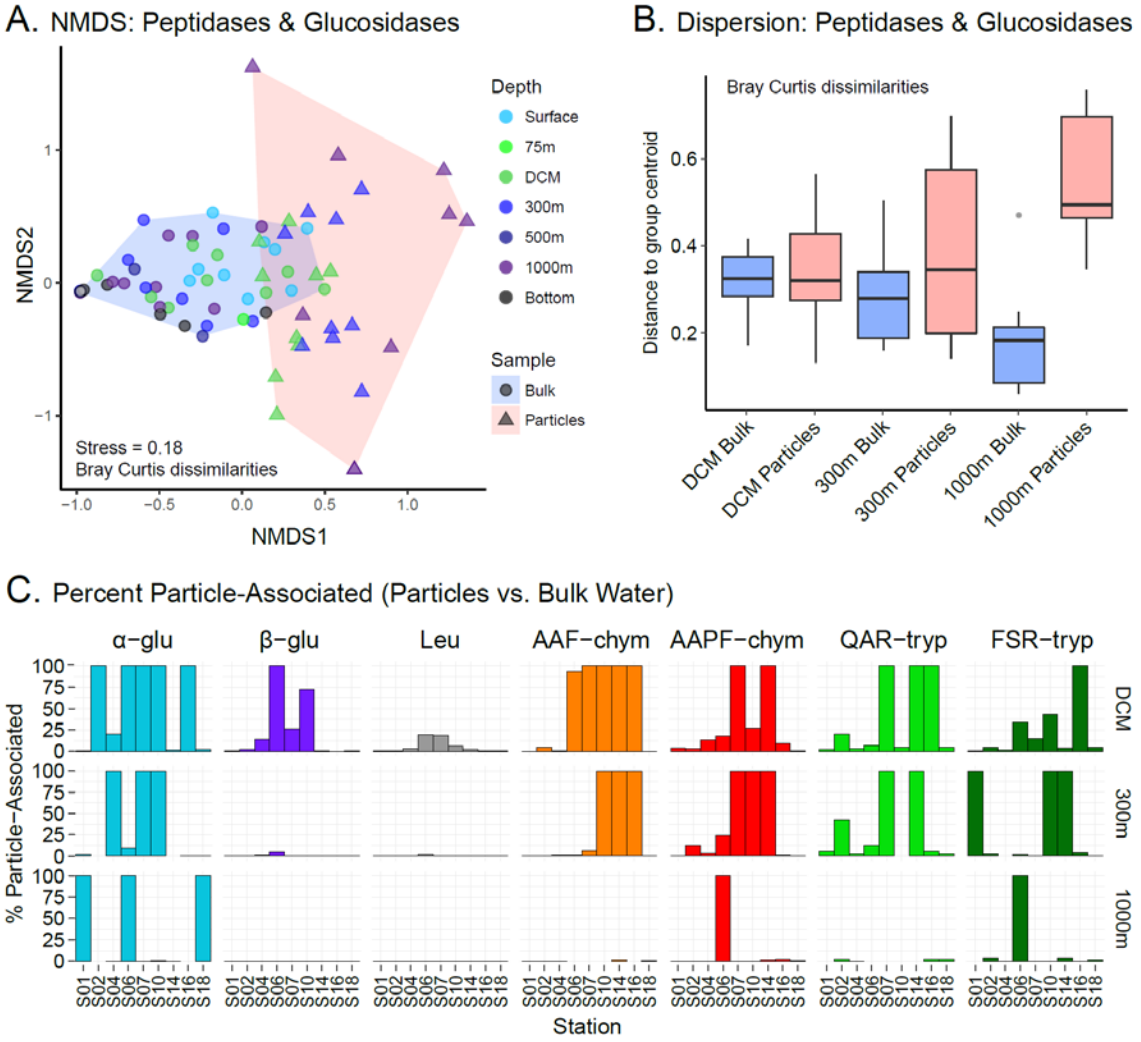


**Figure S4a-c.** Non-metric multidimensional scaling (NMDS) plot of bulk water (blue) and particle-associated (pink) glucosidase and peptidase activities using the Bray-Curtis dissimilarity index (**a**). Bray-Curtis based group dispersions of bulk water and particle-associated glucosidase and peptidase activities, measured as distances of sample rates to group centroid at the DCM, 300 m (mesopelagic), and 1000 m (bathypelagic) (**b**). Proportion of bulk water glucosidase and peptidase activities attributable to the ≥ 3 µm particle-associated fraction (**c**). Note that bulk water peptidase and glucosidase rates used here were measured at the 12 h (t1) timepoint, for comparison to the particle-associated rates at the 24 h (t1) timepoint

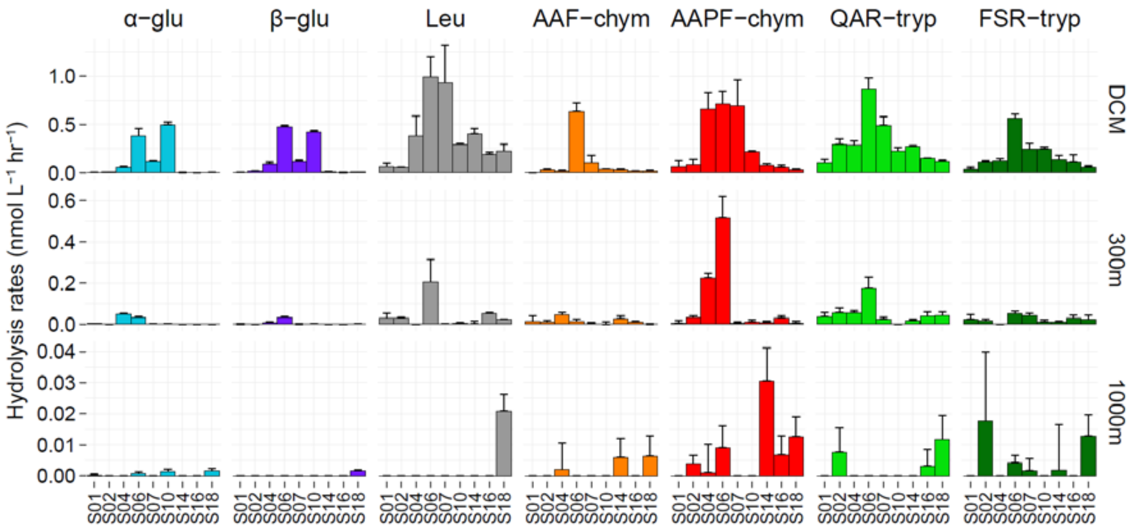


**Figure S5a**. Substrate-separated rates for particle-associated (≥3µm) glucosidase and peptidase activities at the main stations. Note that DCM depths vary by station (*see Table S1*) and y-axis scales differ by depth. Error bars represent the standard deviation of rates from duplicate incubations. Rates shown were measured at 24 h, and normalized by the volume of water filtered for these assays (*see Table S2*). glu = glucosidase, leu = leucine aminopeptidase, chym = chymotrypsin, tryp = trypsin, AAF = alanine-alanine-phenylalanine, AAPF = alanine-alanine-proline-phenylalanine, QAR = glutamine-alanine-arginine, FSR = phenylalanine-serine-arginine.

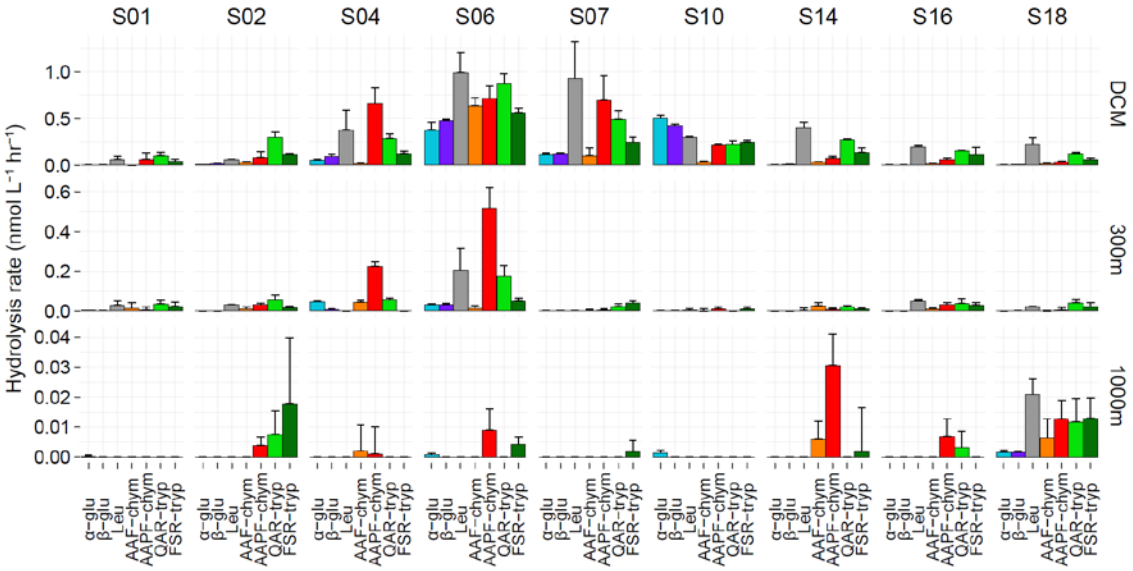


**Figure S5b**. Particle-associated (≥3µm) glucosidase and peptidase activities at the main stations. Note that DCM depths vary by station (*see Table S1*) and y-axis scales differ by depth. Error bars represent the standard deviation of rates from duplicate incubations. Rates shown were measured at 24 h, and normalized by the volume of water filtered for these assays (*see Table S2*). glu = glucosidase, leu = leucine aminopeptidase, chym = chymotrypsin, tryp = trypsin, AAF = alanine-alanine-phenylalanine, AAPF = alanine-alanine-proline-phenylalanine, QAR = glutamine-alanine-arginine, FSR = phenylalanine-serine-arginine.

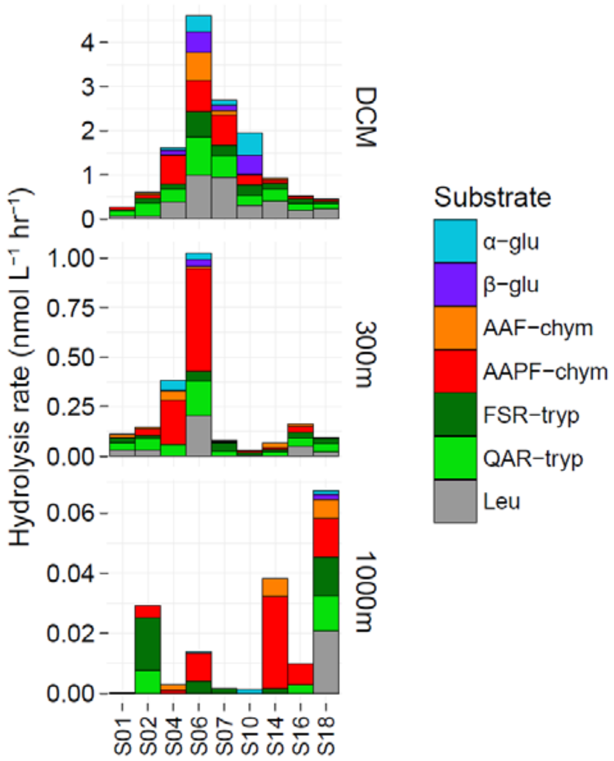


**Figure S5c**. Summed rates of particle-associated (≥3μm) glucosidase and peptidase activities at the main stations. Note that DCM depths vary by station (*see Table S1*) and y-axis scales differ by depth. Error bars represent the standard deviation of rates from duplicate incubations. Rates shown were measured at 24 h, and normalized by the volume of water filtered for these assays (*see Table S2*). glu = glucosidase, leu = leucine aminopeptidase, chym = chymotrypsin, tryp = trypsin, AAF = alanine-alanine-phenylalanine, AAPF = alanine-alanine-proline-phenylalanine, QAR = glutamine-alanine-arginine, FSR = phenylalanine-serine-arginine.

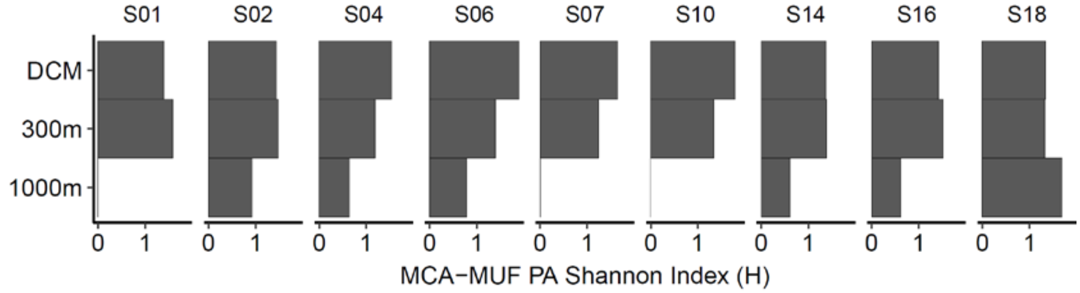


**Figure S5d.** Shannon indices for particle-associated (≥3µm) glucosidase and peptidase activities at all depths at the main stations. Note that DCM and bottom water depths vary by station (*see Table S1*). Shannon indices were calculated from rates measured at 24 h, based on a previously established equation (Steen *et al.*, 2010). N.D. = No Data collected.

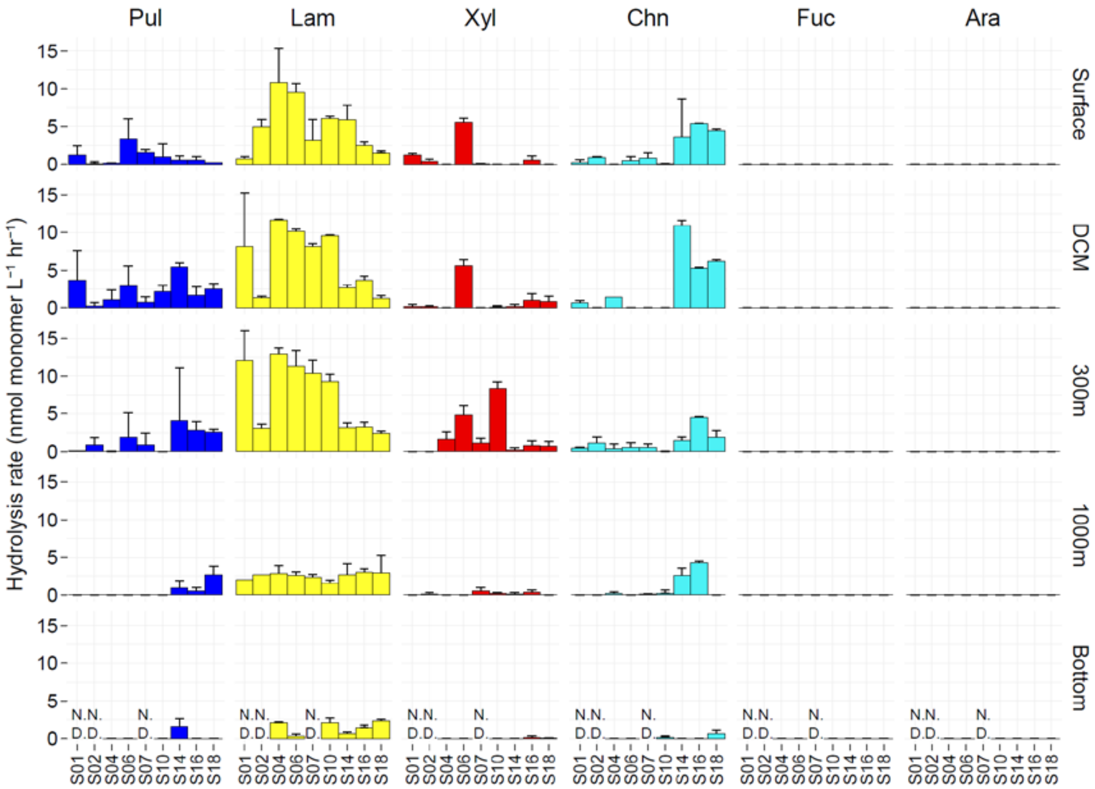


**Figure S6a.** Substrate-separated rates for bulk water (non-size fractionated) polysaccharide hydrolase activities at the main stations. Note that DCM and bottom water depths vary by station (*see Table S1*). Error bars represent the standard deviation of rates from triplicate incubations. Data shown are the maximum rates measured at different timepoints throughout a 25 d incubation period. Pul = Pullulan, Lam = Laminarin, Xyl = Xylan, Chn = Chondroitin Sulfate, Fuc = Fucoidan, Ara = Arabinogalactan, N.D. = No Data collected. There was no detectable hydrolysis for fucoidan and arabinogalactan.

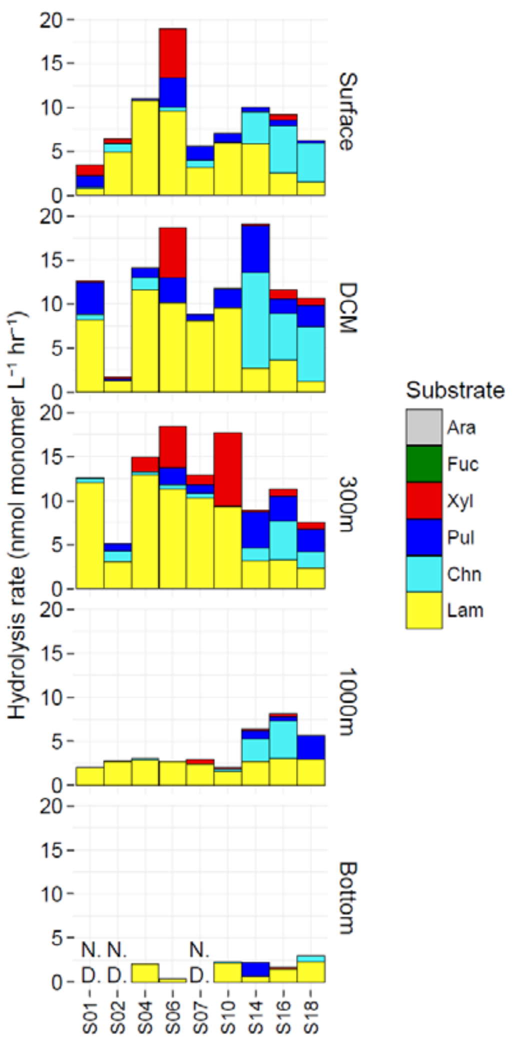


**Figure S6b.** Summed rates for bulk water (non-size fractionated) polysaccharide hydrolase activities at the main stations. Note that DCM and bottom water depths vary by station (*see Table S1*). Error bars represent the standard deviation of rates from triplicate incubations. Data shown are the maximum rates measured at different timepoints throughout a 25 d incubation period. Pul = Pullulan, Lam = Laminarin, Xyl = Xylan, Chn = Chondroitin Sulfate, Fuc = Fucoidan, Ara = Arabinogalactan, N.D. = No Data collected. There was no detectable hydrolysis for fucoidan and arabinogalactan.

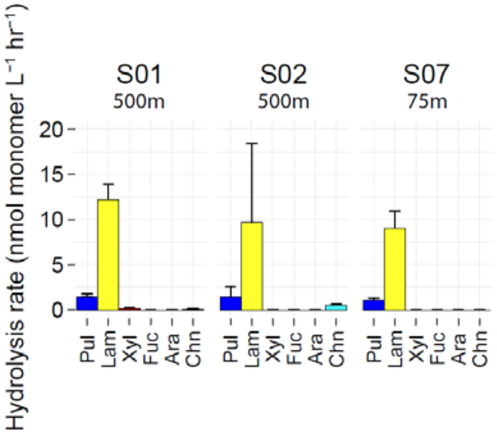


**Figure S6c.** Bulk water (non-size fractionated) polysaccharide hydrolase activities at S01-500m, S02-500m, and S07-75m. Error bars represent the standard deviation of rates from triplicate incubations. Data shown are the maximum rates measured at different timepoints throughout a 25 d incubation period. Pul = Pullulan, Lam = Laminarin, Xyl = Xylan, Chn = Chondroitin Sulfate, Fuc = Fucoidan, Ara = Arabinogalactan.

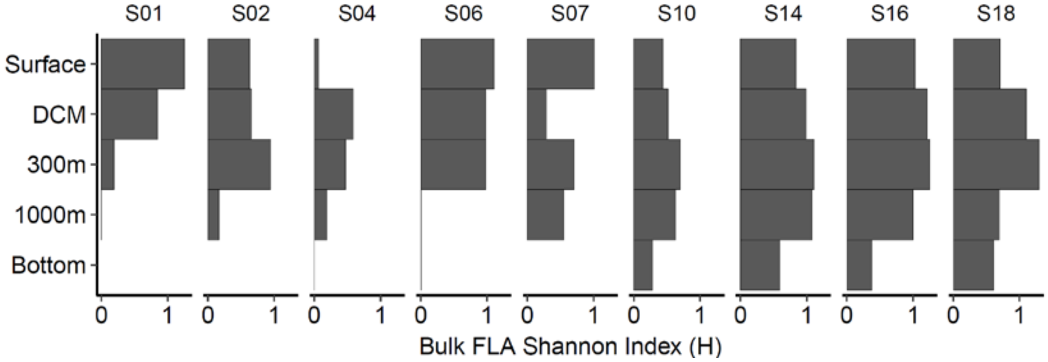


**Figure S6d.** Shannon indices for bulk water (non-size fractionated) polysaccharide hydrolase activities at the main stations. Note that DCM and bottom water depths vary by station (*see Table S1*). Shannon indices were calculated from maximum rates measured at different timepoints, based on a previously established equation (Steen *et al.*, 2010). N.D. = No Data collected.

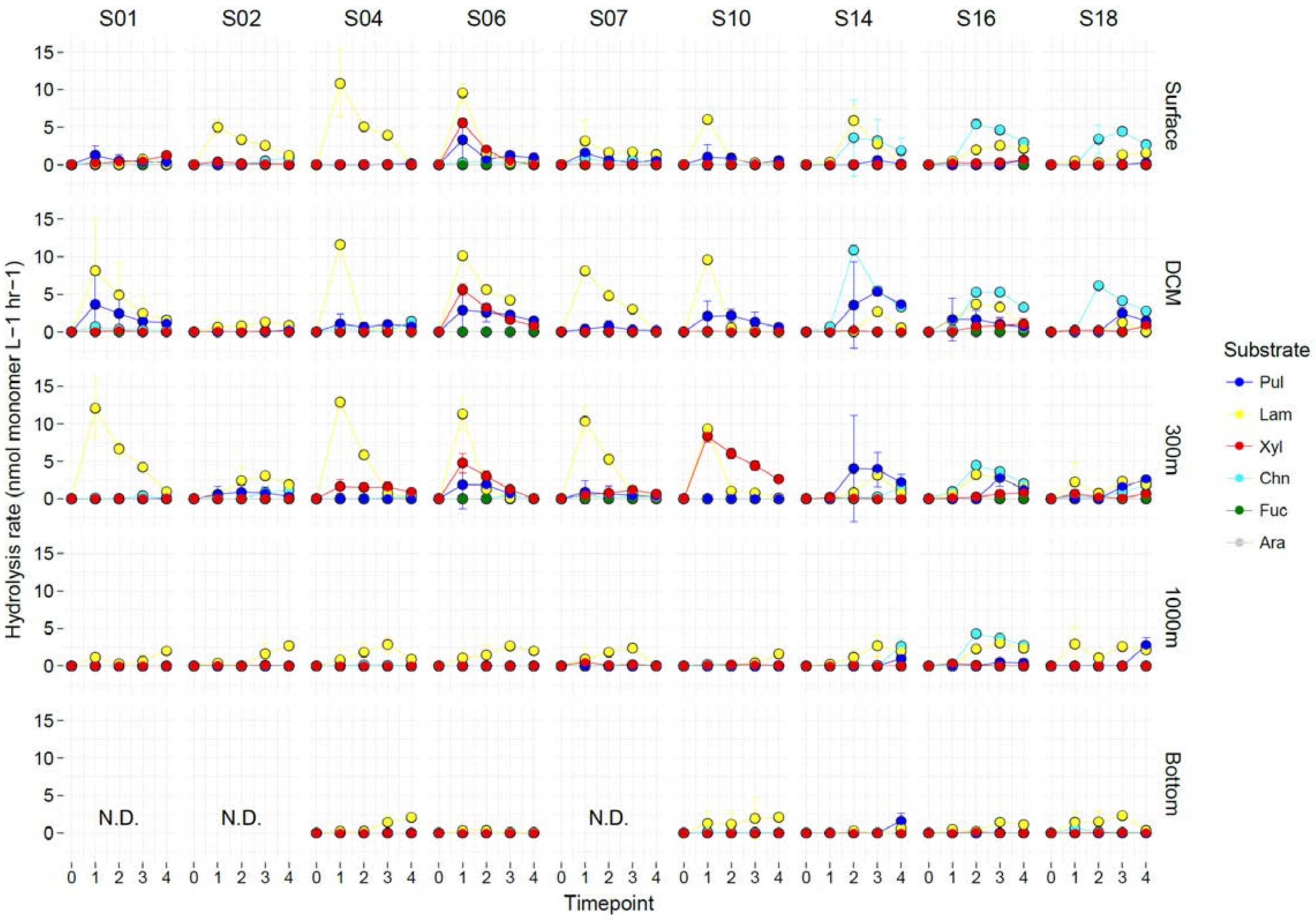


**Figure S6e.** Bulk water (non-size fractionated) polysaccharide hydrolase activities for all timepoints. Note that DCM and bottom water depths vary by station (*see Table S1*). Error bars represent the standard deviation of rates from triplicate incubations. Timepoints 0,1,2,3,4 correspond to 0, 5, 10, 15, 25 days after substrate addition. Pul = Pullulan, Lam = Laminarin, Xyl = Xylan, Chn = Chondroitin Sulfate, Fuc = Fucoidan, Ara = Arabinogalactan.

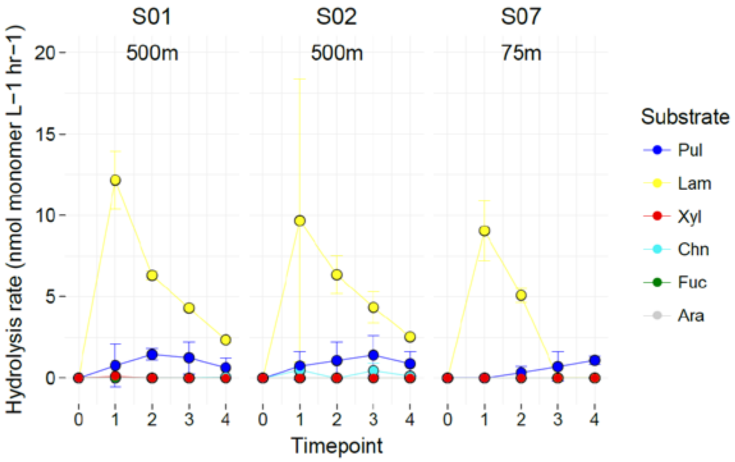


**Figure S6f.** Bulk water (non-size fractionated) polysaccharide hydrolase activities for all timepoints at S01-500m, S02-500m, and S07-75m. Error bars represent the standard deviation of rates from triplicate incubations. Timepoints 0,1,2,3,4 correspond to 0, 5, 10, 15, 25 days after substrate addition. Pul = Pullulan, Lam = Laminarin, Xyl = Xylan, Chn = Chondroitin Sulfate, Fuc = Fucoidan, Ara = Arabinogalactan.

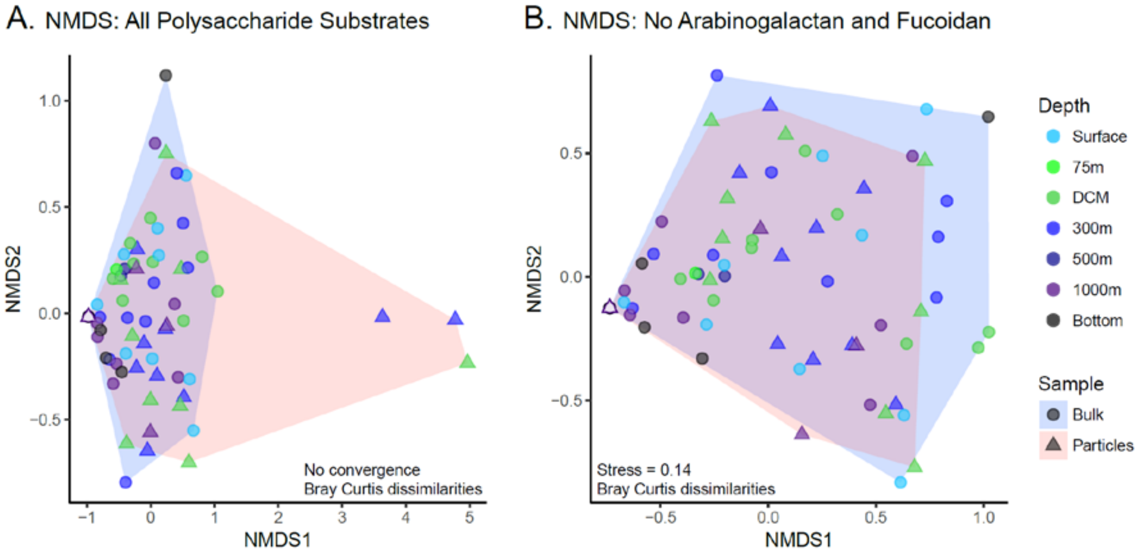


**Figure S7.** Non-metric multidimensional scaling (NMDS) plot of bulk water (blue) and particle-associated (pink) polysaccharide hydrolase activities using the Bray-Curtis dissimilarity index with data from all substrates (**a**) and excluding arabinogalactan and fucoidan (**b**). Because there was no convergence for the NMDS ordination with all substrate data (**a**), data consisting mostly of zeros for arabinogalactan and fucoidan were removed (**b**). Ordination was conducted on maximum rates measured at different timepoints, and with 999 permutations.

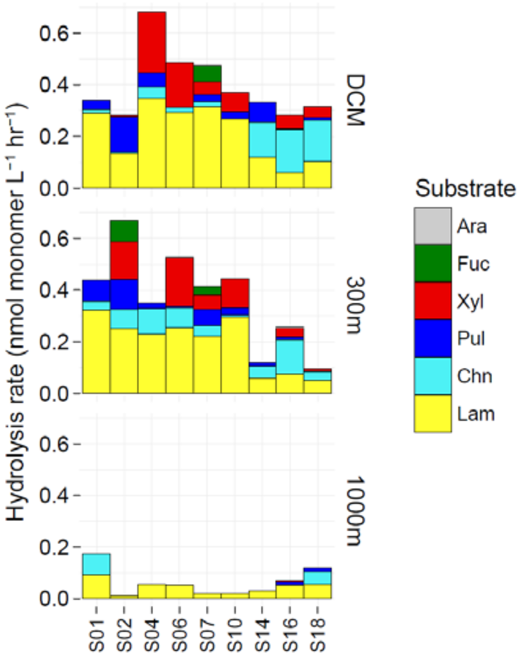


**Figure S7c.** Summed rates for particle-associated (≥3μm) polysaccharide hydrolase activities at the main stations. Note that DCM depths vary by station (*see Table S1*). Error bars represent the standard deviation of rates from triplicate incubations. Data shown are the maximum rates measured at different timepoints throughout a 25 d incubation period, and were normalized by the volume of water filtered for these assays (*see Table S2*). Pul = Pullulan, Lam = Laminarin, Xyl = Xylan, Chn = Chondroitin Sulfate, Fuc = Fucoidan, Ara = Arabinogalactan, N.D. = No Data collected. There was no detectable hydrolysis for fucoidan and arabinogalactan.

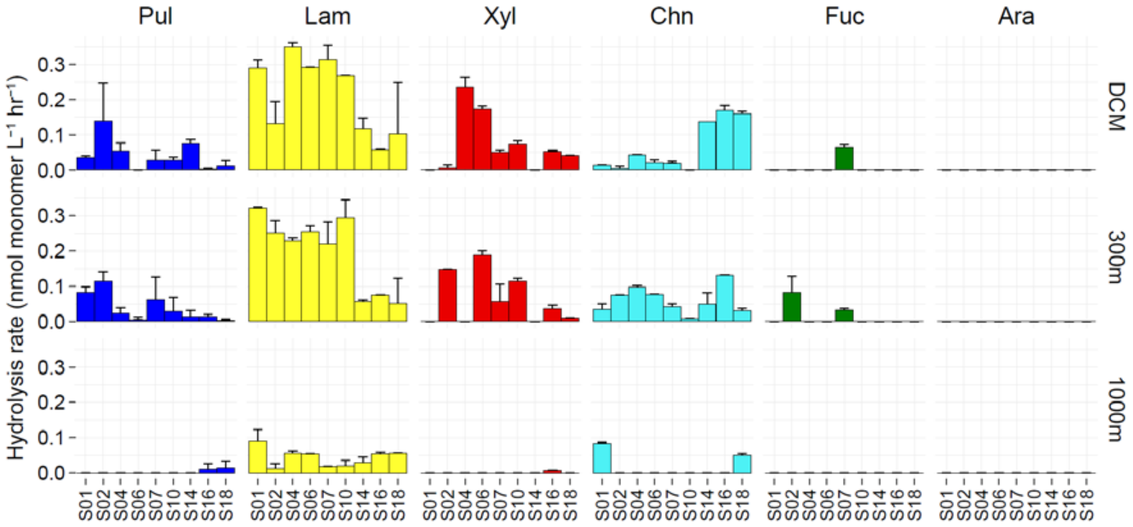


**Figure S7d.** Substrate-separated rates for particle-associated (≥3µm) polysaccharide hydrolase activities at the main stations. Note that DCM depths vary by station (*see Table S1*). Error bars represent the standard deviation of rates from triplicate incubations. Data shown are the maximum rates measured at different timepoints throughout a 25 d incubation period, and were normalized by the volume of water filtered for these assays (*see Table S2*). Pul = Pullulan, Lam = Laminarin, Xyl = Xylan, Chn = Chondroitin Sulfate, Fuc = Fucoidan, Ara = Arabinogalactan, N.D. = No Data collected. There was no detectable hydrolysis for fucoidan and arabinogalactan.

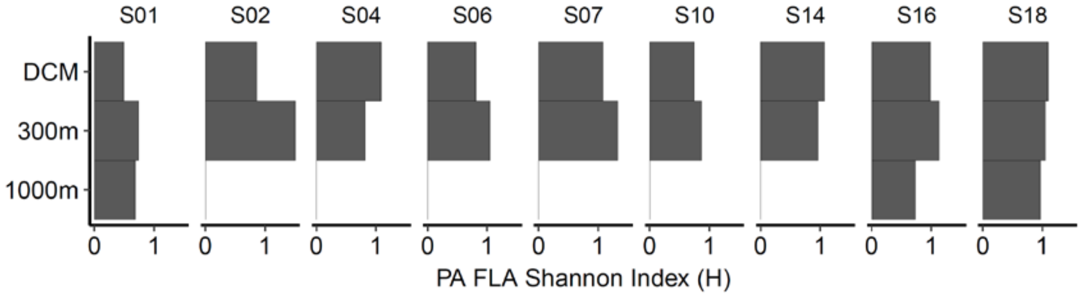


**Figure S7e.** Shannon indices for particle-associated (≥3µm) polysaccharide hydrolase activities at the main stations. Note that DCM depths vary by station (*see Table S1*). Shannon indices were calculated from maximum rates measured at different timepoints, based on a previously established equation (Steen *et al.*, 2010). N.D. = No Data collected.

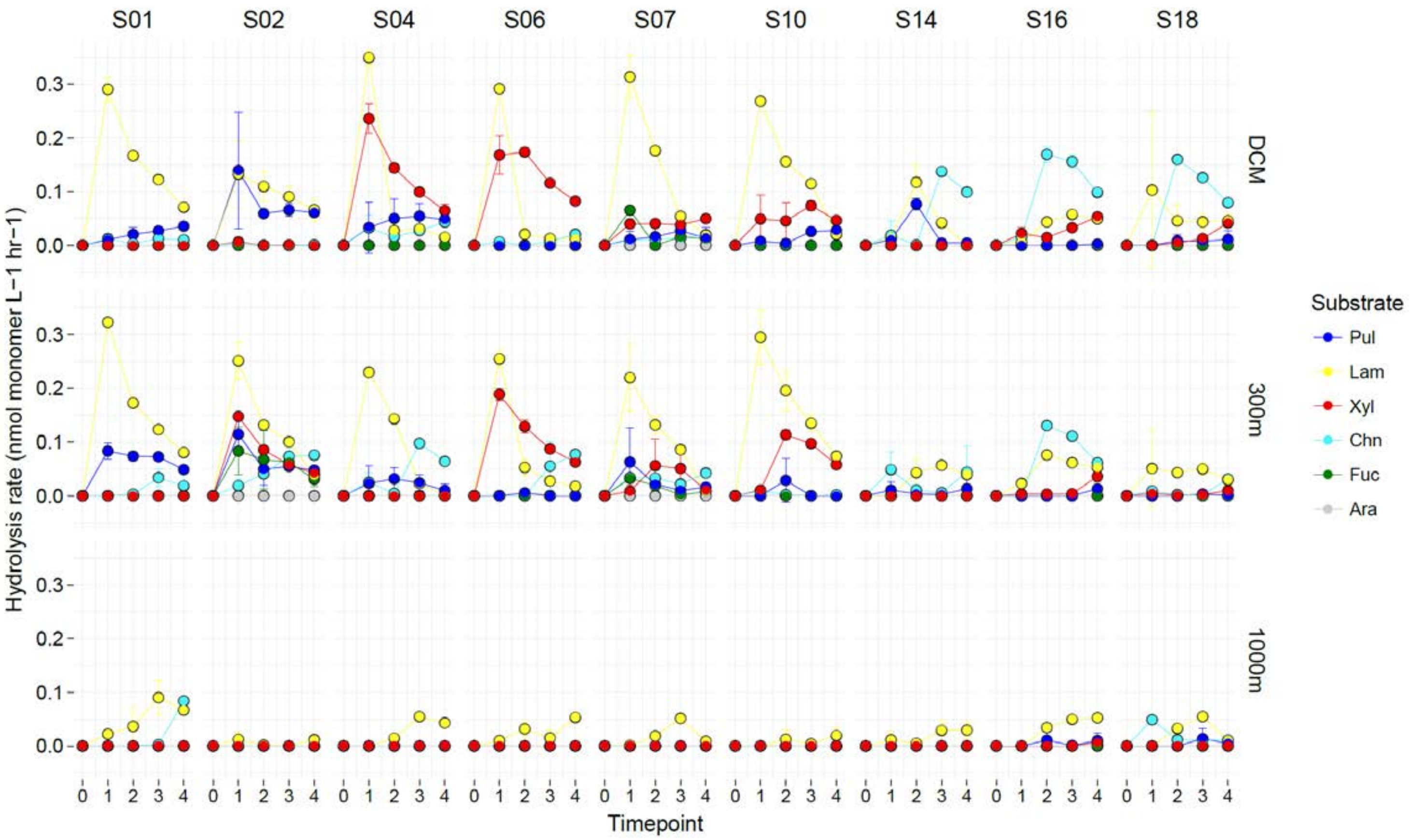


**Figure S7f.** Particle-associated (≥3µm) polysaccharide hydrolase activities for all timepoints. Note that DCM water depths vary by station (*see Table S1*). Error bars represent the standard deviation of rates from triplicate incubations. Timepoints 0,1,2,3,4 correspond to 0, 5, 10, 15, 25 days after substrate addition. Pul = Pullulan, Lam = Laminarin, Xyl = Xylan, Chn = Chondroitin Sulfate, Fuc = Fucoidan, Ara = Arabinogalactan.

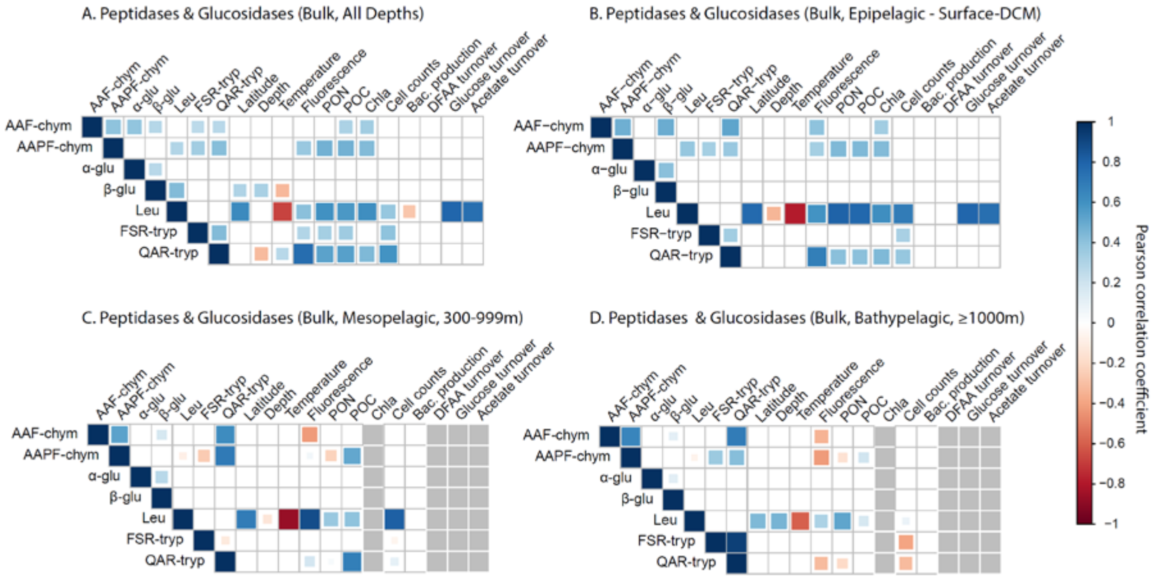


**Figure S8.** Pearson correlations between bulk water (non-size fractionated) peptidases/glucosidases and measured environmental/bulk bacterial activity parameters at all depths (**a**), in the epipelagic (**b**), mesopelagic (**c**) and bathypelagic (**d**). Non-significant correlations ($p < 0.05$) are shown as empty boxes. No rate data due to undetectable enzymatic activities or unavailable environmental/bulk bacterial activity data are shown as gray boxes. Rates analyzed for panels (**a**) and (**b**) include bulk water rates from all stations, whereas those for panels (**c**) and (**d**) are only from the main stations.

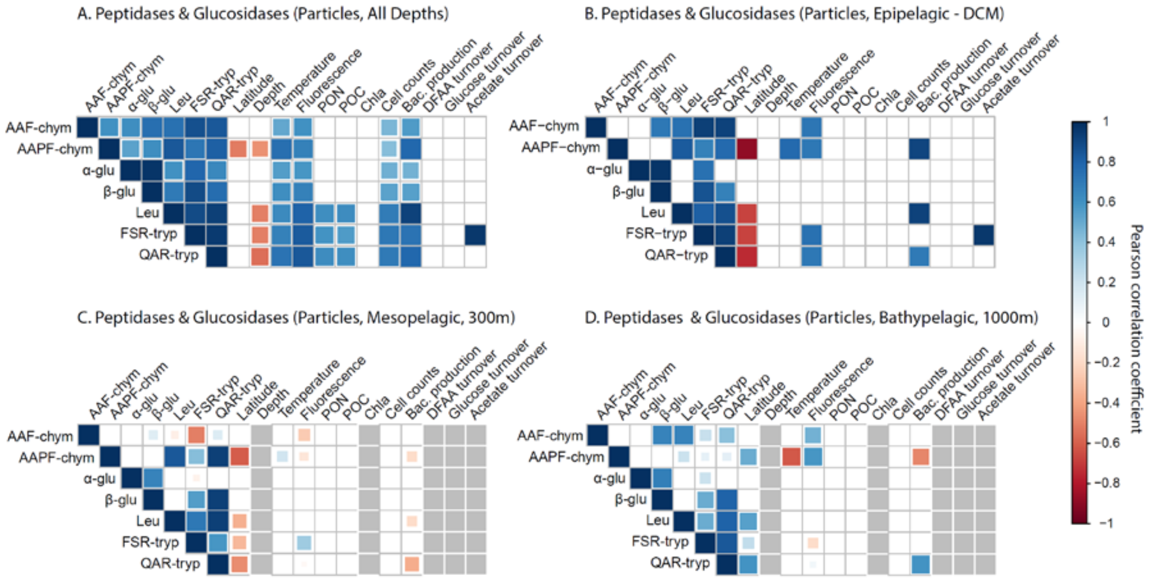


**Figure S9.** Pearson correlations between particle-associated (≥3µm) peptidases/ glucosidases and measured environmental/bulk bacterial activity parameters at all depths (**a**), in the epipelagic (**b**), mesopelagic (**c**) and bathypelagic (**d**). Non-significant correlations ($p < 0.05$) are shown as empty boxes. No rate data due to undetectable enzymatic activities or unavailable environmental/bulk bacterial activity data are shown as gray boxes. Rates analyzed for panels (**a**) and (**b**) include bulk water rates from all stations, whereas those for panels (**c**) and (**d**) are only from the main stations.

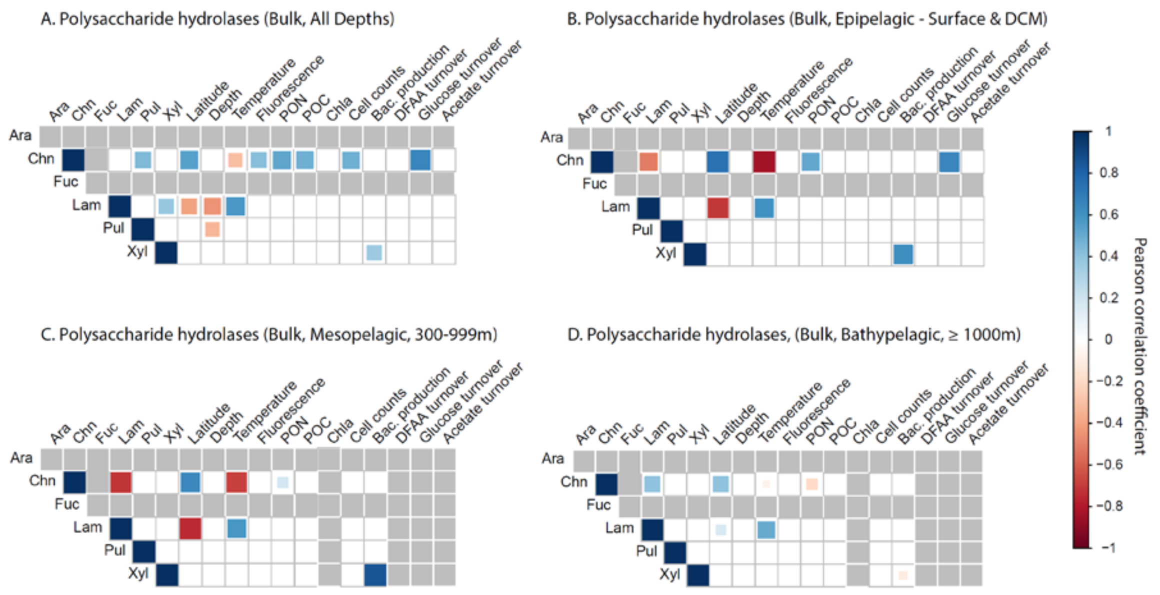


**Figure S10.** Pearson correlations between bulk water (non-size fractionated) polysaccharide hydrolases and measured environmental/bulk bacterial activity parameters at all depths (**a**), in the epipelagic (**b**), mesopelagic (**c**) and bathypelagic (**d**). Non-significant correlations ($p < 0.05$) are shown as empty boxes. No rate data due to undetectable enzymatic activities or unavailable environmental/bulk bacterial activity data are shown as gray boxes. Rates analyzed for panels (**a**) and (**b**) include bulk water rates from all stations, whereas those for panels (**c**) and (**d**) are only from the main stations.

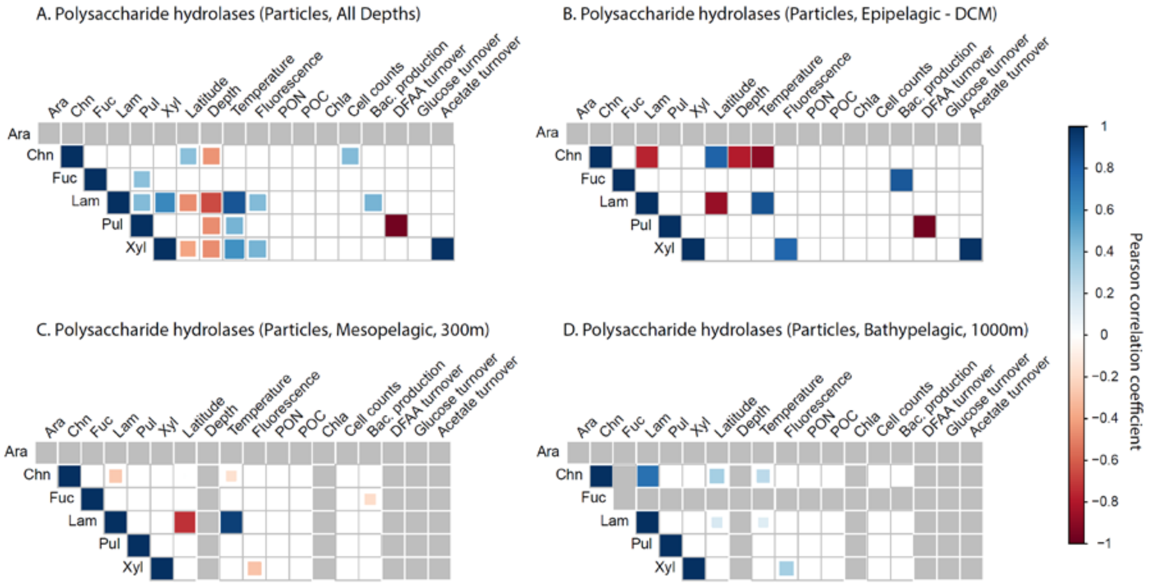


**Figure S11.** Pearson correlations between particle-associated (≥3μm) polysaccharide hydrolases and measured environmental/bulk bacterial activity parameters at all depths (**a**), in the epipelagic (**b**), mesopelagic (**c**) and bathypelagic (**d**). Non-significant correlations ($p < 0.05$) are shown as empty boxes. No rate data due to undetectable enzymatic activities or unavailable environmental/bulk bacterial activity data are shown as gray boxes. Rates analyzed for panels (**a**) and (**b**) include bulk water rates from all stations, whereas those for panels (**c**) and (**d**) are only from the main stations.
